# The Cretan Carob Virome

**DOI:** 10.64898/2026.09.15.751538

**Authors:** Konstantina Katsarou, Maria-Eleni Biti, Androniki Foteinou, George Papagiannakis, Despina Michelaki, Christos Andronis, Grigorios Emvalomatis, Kriton Kalantidis

## Abstract

Carob (*Ceratonia siliqua*) is an important yet often underestimated Mediterranean tree species. Historically used as a sweetener by ancient Greeks and Arabs, it also supported populations during periods of famine. Beyond its cultural value, carob exhibits notable agronomic traits, including drought and temperature tolerance, soil-erosion prevention, and potential contributions to rural development. Although a Cretan carob genome assembly has recently become available, little is known about pathogens associated with this species, particularly viruses. Here, we characterized the virome of carob trees in Crete using high-throughput sequencing (HTS) and targeted molecular validation. We collected 134 leaf samples from 34 growers across the island’s four prefectures. HTS analysis revealed novel isolates of two *Secoviridae* members (Artichoke yellow ringspot virus - AYRSV) and Strawberry latent ringspot virus - SLRSV), as well as Soybean ilarvirus 1 (SolV1), a putative and incompletely characterized jivivirus, and a previously undescribed coguvirus of the family *Phenuiviridae*, all confirmed by PCR screening. Phylogenetic and sequence analyses support the classification of the coguvirus as a distinct viral species, for which we propose the name Carob coguvirus 1 (CaCoV1). To our knowledge, this study provides the first comprehensive characterization of the carob virome. These findings expand current knowledge of viral diversity in carob and establish a foundation for future epidemiological studies, certification of propagation material, and the sustainable expansion of carob cultivation in Crete and other Mediterranean regions.

**IMPACT STATEMENT:** This study provides the first comprehensive characterization of the virome associated with carob trees. It reveals previously unknown viral diversity, including a proposed novel coguvirus, and identifies viruses that may affect plant health and the production of propagation material. The results provide a basis for future virus-surveillance programs, improved propagation and grafting practices, and more resilient carob cultivation, supporting ongoing efforts to revitalize this important Mediterranean crop.

## INTRODUCTION

*Ceratonia siliqua (C. siliqua),* or carob tree, is a member of the Fabaceae family and is mostly found in Mediterranean countries, like Spain, Portugal, Greece, Turkey, Cyprus, Morocco, and Syria [1]. The name of the tree comes from the Greek word ‘*Keras*’, which means horn, and the Latin word ‘*siliqua*’, alluding to the hardness and shape of the pod. Carob trees were probably abundant in the Middle East in ancient times. Both the ancient Greeks and Arabs recognized the value of this tree and spread it along the Mediterranean and North African coasts. Nowadays, these trees can also be found in regions with similar climate conditions, like California, Mexico, Australia, South Africa, and India [1].

Carob trees can reach heights of up to10m, do not shed leaves in autumn, but rather in July every second year, and they only renew leaves in April and May. Carob is extremely well adapted to drought and can tolerate temperatures of −6 °C to +50 °C. It is an excellent tree to use for preventing soil erosion and has the potential to contribute to the development of rural areas. In the past few years, the carob industry has been developing in multiple directions, with leaves, pods, and seeds being used to produce a variety of products, each with its own advantages [1, 2]. Carob products are rich in sucrose and protein, but poor in fat. Therefore, they are considered an excellent food source for farm animals, children, or people [1]. Carob is increasingly recognized as a crop with significant health-related potential, exhibiting anti-cardiovascular, antioxidant, and gastrointestinal benefits [3].

In terms of cultivation practices, carob trees require little care, which includes pruning, weeding, and, of course, harvesting, and little to no fertilization or irrigation. Although production involves limited amounts of materials, the forgone profit from alternative use of the occupied land area (opportunity cost) is non-negligible, as the time from planting until first harvesting is six to eight years, while full productive maturity is achieved 20-25 years after planting [1]. However, once a carob tree is ready to be harvested, it can remain productive for up to 100 years. Traditionally, carob trees on the island of Crete are grown either in specialized orchards or in the same fields with other trees, as a side crop in multi-cropping systems. Including the labor-intensive activities of harvesting, whether hired or own labor is used, it is not clear whether carob production at a commercial scale is economically viable in the region, while continuation of its cultivation faces challenges stemming from alternative crops and the abandonment of farming [4].

Carob production can be influenced by many biotic factors, especially pathogens and pests (P&P). Yet, despite the long history of carob cultivation in the Mediterranean, we still know surprisingly little about the organisms that affect its health. Only a handful of fungal pathogens have been described so far, including *Pseudocercospora ceratoniae*, *Alternaria alternata*, and *Pestalotiopsis uvicola* [5–8]. Insects are far more diverse: surveys across Mediterranean countries have recorded thirty-one species (mostly beetles, weevils, and moths) living on carob trees. Even so, the actual damage they cause remains poorly documented, and in many cases completely unknown [8, 9]. Bacterial infections have also been mentioned, such as *Pseudomonas syringae* pv. *ciccaronei*, but no recent work has confirmed its presence or clarified its impact [10]. Finally, small mammals like rodents or goats can damage the plants and have been associated with large economic losses [8].

However, to our knowledge, there is no information available about infections with other pathogens and particularly viruses, well-known contributors to yield loss in many perennial crops. In this work, we investigated the virome of Cretan carob trees. We collected leaf tissue samples from all four prefectures of the island of Crete in Greece, performed high-throughput sequencing, and identified new isolates of known viruses either already present or identified for the first time in Greece as well as completely new viruses.

## Material and Methods

### Tissue collections

Leaf tissue samples were collected from all four prefectures of the island of Crete in the fall of 2024. Detailed collection dates and coded names used throughout this project are presented in Table S1. Upon arrival at the laboratory, all samples were photographed and stored at −80°C. All plant tissue samples were collected from growers with their prior written consent, and the study was conducted under the approval of the University’s Bioethics Committee (23/09.02.2024).

### RNA extractions and High-throughput sequencing

After testing various protocols, we opted for RNA extraction with the Monarch® Total RNA Miniprep Kit (New England Biolabs, Ipswich, MA, USA) according to manufacturer instructions, including a DNAseI treatment step as advised in the kit. 350 ng of each sample were loaded in a 1% agarose/formaldehyde gel as has been described before, to assess their quality (Figure S1). Only high-quality RNAs were used for further analysis. Samples were pooled by combining 10 or 11 individual samples, contributing 350 ng from each. In total, 13 pooled samples were used for high-throughput sequencing (HTS) analysis in Macrogen Europe (Amsterdam, The Netherlands). Libraries were constructed using the Total RNA Library Prep Plant Kit with RiboZero, and paired-end reads of 151 nt were generated using the Illumina NovaSeq Platform. The numbers of reads obtained per sample is presented in Table S2.

### Bioinformatic analysis to identify viruses

Bioinformatic analysis was performed as previously described [11, 12]. In brief, raw read quality was assessed with FastQC [13], and adapters were trimmed with fastp [14] (-q 20, --length_required 21, --cut_tail, --cut_front, --cut_mean_quality 20). rRNA sequences were removed using BBDuk (ribokmers.fa; default settings) (https://sourceforge.net/projects/bbmap/) [15]. Host reads were filtered by aligning to the carob genome recently published by our laboratory [16] using BBSplit and the remaining reads were *de novo* assembled using SPAdes (-meta) [15, 17]. Contigs >200 nt were analysed by BLASTn (NCBI Blast+ v2.9.0) [18] against RVDB v29.0 [19, 20], and taxonomic assignments were added using taxonomizr (https://cran.r-project.org/web/packages/taxonomizr/index.html).

### Validation of viral presence

Complementary DNA (cDNA) was synthesized using 500 ng of total RNA, with M-MuLV reverse transcriptase and random hexamers obtained by EnzyQuest (Heraklion, Crete, Greece) using the manufacturer’s proposed protocol. The cDNA was diluted 1:10 in nuclease-free water, and PCR was performed using Taq DNA polymerase and virus-specific primers again from EnzyQuest. PCR products were visualized on 1.5 % agarose gels, excised, purified with NucleoSpin Gel and PCR Clean-Up (Macherey-Nagel, Germany), and Sanger-sequenced by Genewiz (Leipzig, Germany) or Macrogen Europe (Amsterdam, The Netherlands). For full-genome amplification of putative novel viruses, primers were designed from HTS data, and overlapping fragments were amplified from cDNA generated as above. All used primers and Tms are listed in Table S3.

### Phylogenetic analyses

Phylogenetic analyses were conducted using both nucleotide (nt) and amino acid (aa) sequence datasets, depending on the genomic region, virus examined, and data availability. A standardized pipeline was established on the GalaxyEU platform and MEGA X [21] software as follows: sequences retrieved from NCBI were aligned using MAFFT [22] and the optimal substitution model was determined in MEGA X. Maximum Likelihood (ML) trees were generated with IQ-TREE [23] (1,000 bootstrap replicates) and visualized in iTOL[24] excluding bootstrap values below 75%. For Artichoke yellow ringspot virus (AYRSV), intraspecies phylogenetic analysis was performed at the nucleotide level using partial RdRp sequences, specifically motif A and the upstream region, using the best-fit substitution model K80 + I, as reported by Karapetsi *et al* [25]. For Strawberry latent ringspot virus (SLRSV), intraspecies phylogenetic analyses were performed at both nt and aa levels using the ProPol region between the CG motif of the proteinase and the GDD motif of the polymerase and CPs, both large and small proteins. The best-fit models were JTT + G + I for amino acid and GTR + G + I for nucleotide analyses. Next, the intragenus phylogenetic analysis for Soybean ilarvirus 1 (SoIV1) was limited to the MP and CP proteins due to insufficient ProPol data from the SoIV1 carob isolate. The best-fit models were LG + G + I for MP and LG + G + I + F for CP. Finally, for the newly identified Coguvirus, amino acid-level phylogenetic analyses were performed for the L (both full and core sequence), N, and MP proteins. Representative sequences from each known coguvirus were included, along with cogu-like Bocivirus sequences. The best-fit evolutionary models were LG + G, LG + G + I + F, and JTT + G + I, respectively. The IDs of all used sequences are noted in the figures.

### Softwares

Photoshop CS6, was used to assemble pictures. Mapchart was used to create the map of Crete (https://www.mapchart.net/). Geneious version 7.1 (Biomatters - http://www.geneious.com) was used for alignments. NCBI primer 3 (https://www.ncbi.nlm.nih.gov/tools/primer-blast/index.cgi?ORGANISM=9913&INPUT_SEQUENCE=NM_001443850.1) was used for primer design. ORFFinder was used for identification of ORFs in the obtained sequences (https://www.ncbi.nlm.nih.gov/orffinder/) and CD-Search for identification of specific motifs within the obtained proteins (https://www.ncbi.nlm.nih.gov/Structure/cdd/wrpsb.cgi) [26]. The Vienna RNA Websuite was used for RNA structures [27]. The Weblogo software was used for visualization in supplementary Figure S13 (https://weblogo.threeplusone.com) [28]. Finally, SDTv1.3 was used to analyze sequence similarity [29].

## Results - Discussion

### Estimation of carob production in Crete

According to the Food and Agriculture Organization of the United Nations (FAO), which has recorded production data since 1961, Spain and Portugal are the largest carob producers, with a total production of 36.4 thousand tonnes (t) and 41.3K t respectively, followed by Italy (28,9K t), Morocco (22,2K t), Greece (12,3K t), Türkiye (15K t), and Cyprus (12,8K t)^1^. In Greece, approximately 80 % of carob production is concentrated on the island of Crete. Giving data from the Hellenic Statistical Authority, carob production on Crete is primarily carried out under two production systems:(1) as a monoculture, in which the trees are relatively young (<20 years) and (ii) in combination with the production of other perennial crops, where the trees are scattered in the groves, mainly of olive and, less frequently, of citrus trees or with annual crops, which are cultivated mainly to be used as animal feed. The monoculture system is relatively new in the region and it was introduced through subsidies supporting the establishment of new carob orchards on land of marginal productivity, as well as through renewed interest in the production of carob products and byproducts for human consumption. However, these measures/developments did not have a significant impact on overall production, and were largely offset by abandonment of orchards due to the expansion of alternative economic activities, such as tourism and the preference of cultivators for other perennial crops, such as olives or grapevines (Figure 1). Thus, a gradual reduction in the number of trees has been observed over the past two decades^2^, with a minor recovery in recent years. Total production follows a similar trend, with large increases appearing in years when the selling price of carob pods is large enough to cover the cost of harvesting. The mixed production system is a low-intensity system in which labor is the primary variable input, used mostly during harvesting and, infrequently, for pruning the trees and weeding. In this traditional system, carob production is not the farm’s primary output, and harvesting may not take place when the prevailing farm-gate price is too low to cover labor costs.

**Figure 1:**
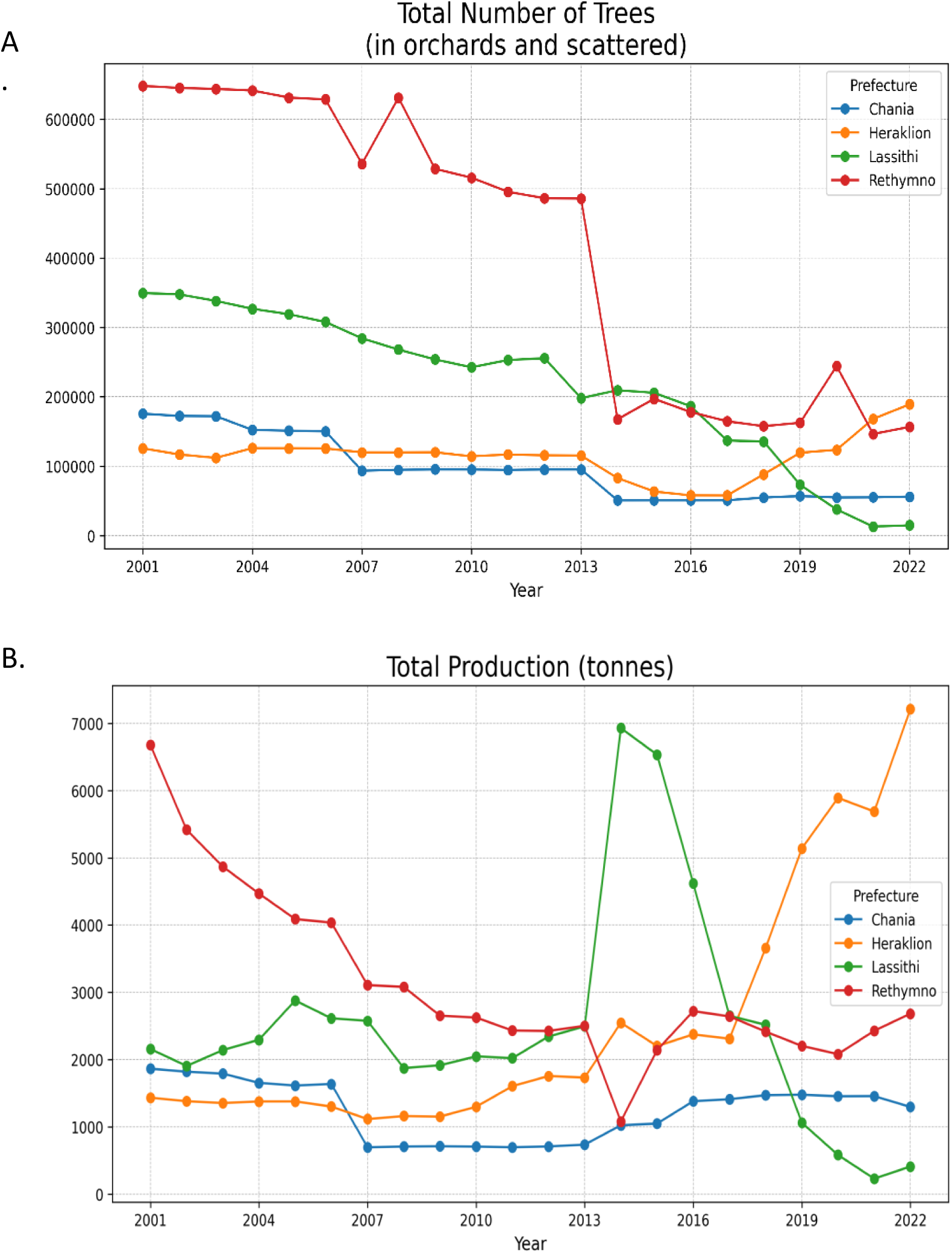
Evolution of carob production by prefecture on the island of Crete. A) Total number of trees, B) Total carob production (in tonnes). Source: Hellenic Statistical Authority, tailor-made statistical data provided upon request.

In this work, we identified 34 different carob producers in the four prefectures of the island (Chania, Rethymnon, Lasithi and Heraklion) (Figure 2). The data collection took place in two stages. Information on the production system and the characteristics of farms and farm operators was collected through face-to-face interviews conducted from the beginning of the production period in late August until the middle of the production period in December. At this stage, respondents were also asked to provide an estimate of their production volume for the coming year. Actual data on output and variable inputs were collected and combined midway through the production year and after harvesting. Table 1 presents summary statistics on key variables regarding the farm and farm operator characteristics. 80 % of the farms used the mixed production system, and the median number of trees was 85, while there were a few farms (using the monoculture system) with more than 1000 trees. Farm operators were relatively old (mean and median age approximately 60 years), and only 35 % of the growers made farming their main occupation. Furthermore, the 2024-25 production year was marked by unfavorable weather conditions. Dry weather occurred in many parts of the island during the flowering period, from August to October, and during pod development, from April to June 2025. These conditions reduced yields in some areas to such an extent that harvesting carob pods was not economically viable for some growers. In addition, the difference between expected harvest, as stated by the farmers in late 2024, and realized harvest, as stated in 2025, was very large, reaching 49% in the sample.

**Figure 2:**
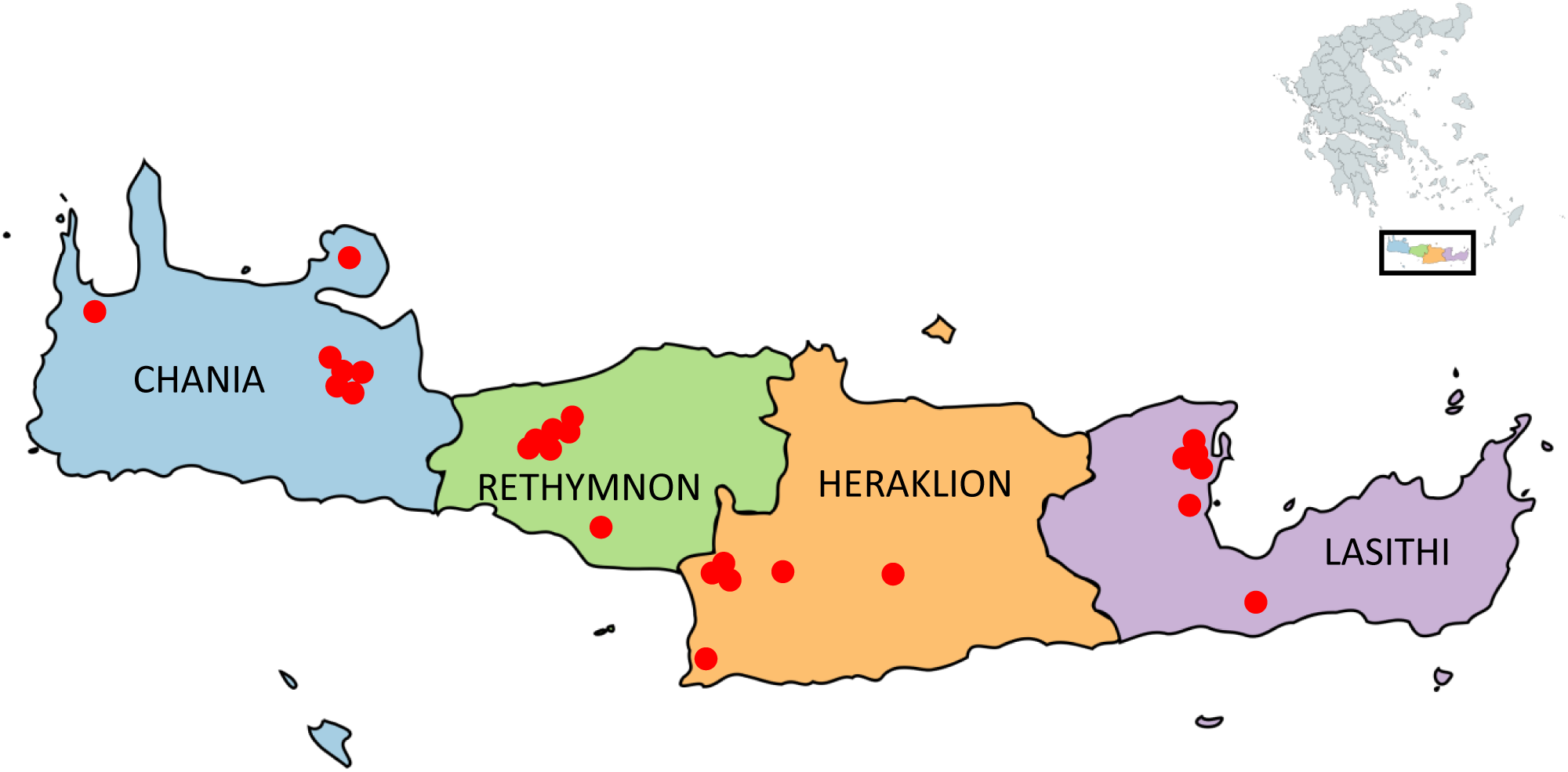
Geographical distribution of the sample collection across all four regional units of the island of Crete

**Table 1:** Information on the 34 Carob cultivars and the trees sampled in this study.

| **Variable** | **Mean / Count** | **Median / Percentage (%)** | **Std. Dev.** |
| --- | --- | --- | --- |
| **Production System** |  |  |  |
| Mixed system | 27 | 79.4% |  |
| Monoculture | 7 | 20.6% |  |
| **Number of Trees** | 270.15 | 85 | 490.6 |
| **Age** | 59.41 | 60.5 | 10.82 |
| **Operator's Main Occupation** |  |  |  |
| Farming | 12 | 35.3% |  |
| Other | 22 | 64.7% |  |
| **Education** |  |  |  |
| Incomplete primary education | 2 | 5.9% |  |
| Complete primary education | 4 | 11.8% |  |
| Completed 9-year education | 8 | 23.5% |  |
| Completed secondary education | 5 | 14.7% |  |
| University degree | 11 | 32.4% |  |
| Post-graduate degree | 4 | 11.8% |  |

Taken together, our analysis indicates that carob production in Crete relies largely on old trees, which are often passed down through generations, and therefore remains a secondary source of income. Nevertheless, it could become a profitable activity for new growers if market prices continue to rise and if interest in carob cultivation increases.

### Identification of viruses in carob trees

We next investigated the health condition of these (mostly old) carob trees existing in Crete. For this, we collected leaf tissue samples from the identified 34 producers from September to December of 2024 (Figure 2). Sampling focused primarily on trees with possible symptoms. We identified a variety of symptoms, some of which resembled those previously described in other studies as being potentially associated with fungal or bacterial infections, and also detected a few insects (Figure S2). In total, we collected 134 samples, and a unique coded name was attributed to each one of them (Table S1). RNA was extracted, and the quality was assessed on a denaturing gel (Figure S1). Samples were then pooled (by 10 or 11), and high-throughput sequencing (HTS) was performed. In total, 13 pooled samples were sequenced, and the analysis revealed a range of viruses from different families, which are described in the following sections.

### Family *Secoviridae*

*Secoviridae* is a family of non-enveloped viruses containing either a monopartite or a bipartite linear positive-sense RNA with a size ranging from 9 to 13.7 kbp, enclosed in particles of 25 to 30 nm [30]. The family includes one sub-family with four genera, six additional genera, three sub-genera, and 165 species according to the International Committee on Taxonomy of Viruses (ICTV) (https://ictv.global/report/chapter/secoviridae/secoviridae).

#### Artichoke yellow ringspot virus

Artichoke yellow ringspot virus (AYRSV) (species *Nepovirus cynarae*) belongs to the Nepovirus genus, containing viruses with bipartite genomes. RNA1 has a size of approximately 7 Kb and contains replication-associated proteins, including RNA-dependent RNA polymerase (RdRp), and RNA2 is about 6 Kb coding coat and movement proteins. The virus is usually transmitted by nematodes, however, seed and pollen transmission have also been reported. AYRSV causes ringspot symptoms, malformations and/or tip necrosis and has been found to infect plants from *Amaranthaceae*, *Asteraceae*, *Caryophyllaceae*, *Cucurbitaceae*, *Fabaceae*, *Lamiaceae*, *Nyctaginaceae,* and *Solanaceae* families [25, 31–36]. This virus has circulated mainly in Greece, Italy, and Turkey since the 1970s, but its ability to infect economically important crops makes its presence in other countries very likely.

Among the 13 pools we have analyzed by HTS, we identified 55 contigs of the virus with sizes from 207 nt to 6753 nt. As shown in Table S4, these contigs exhibit sequence similarity ranging from 84.08 % to 97.99 %. Following, we performed PCR using specific primers (Table S3) in all samples to identify trees containing the virus (Figure S3 - Table S5). Out of the 134 carob trees analyzed, we identified 12 virus-positive samples (detection frequency of 8.95 %). Two samples from Chania region were Sanger sequenced (Δ2 and Σ4) and showed a 99,6 % and 92,20 % compared with the longer node identified in the respective pools, suggesting the circulation of distinct viral variants. The highest incidence of AYRSV-positive trees was observed in the Chania regional unit, where 8 of the 12 positive samples were detected. This distribution suggests that viral circulation may be more extensive in that area.

To identify a possible phenotype, we examined the available photographs. We observed symptoms that ranged from mild yellowing of the leaves to more pronounced phenotypes like reddish ringspots (Figure 3, Figure S4, Table S5). We also observed tip necrosis in branchlets, but not necessarily along the entire branch, resulting in a peculiar phenotype in which branches bore both necrotic and fully viable leaves (Figure 3, Figure S4). To investigate whether the detected AYRSV isolates could be mechanically transmitted to experimental hosts, we inoculated *Phaseolus vulgaris* and *Nicotiana benthamiana* plants with sap extracted from AYRSV-positive carob leaves. However, no infections were obtained, suggesting that either the inoculation method was not appropriate or that these particular viral strains do not infect the chosen plant species.

**Figure 3:**
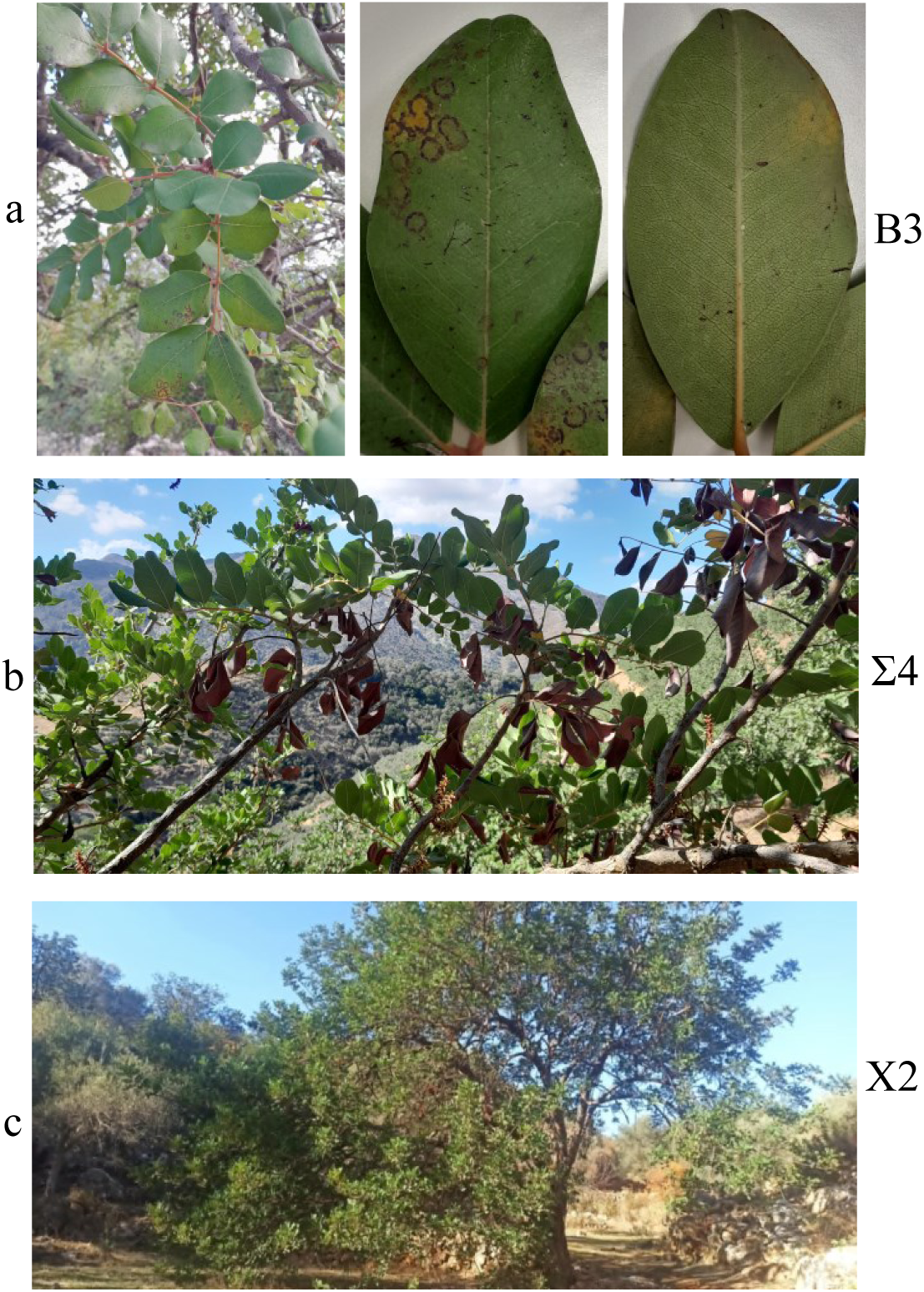
Representative phenotypes observed in AYRSV-infected carob trees from three different trees a) B3, b) Σ4 and c) X2.

We then performed a phylogenetic analysis of a part of the PRO-POL combined with a region of 100 bp upstream according to Karapetsi *et al* [25]. As the authors note, three subgroups of AYRSV have been identified. In Figure 4, we show that the AYRSV isolates identified in Cretan carob trees were part of Subgroup III. Furthermore, these isolates are closely related to a strain isolated from *Vicia faba* in the region of Crete (AM087672.1), suggesting that these isolates are circulating in various plants on the island of Crete.

**Figure 4:**
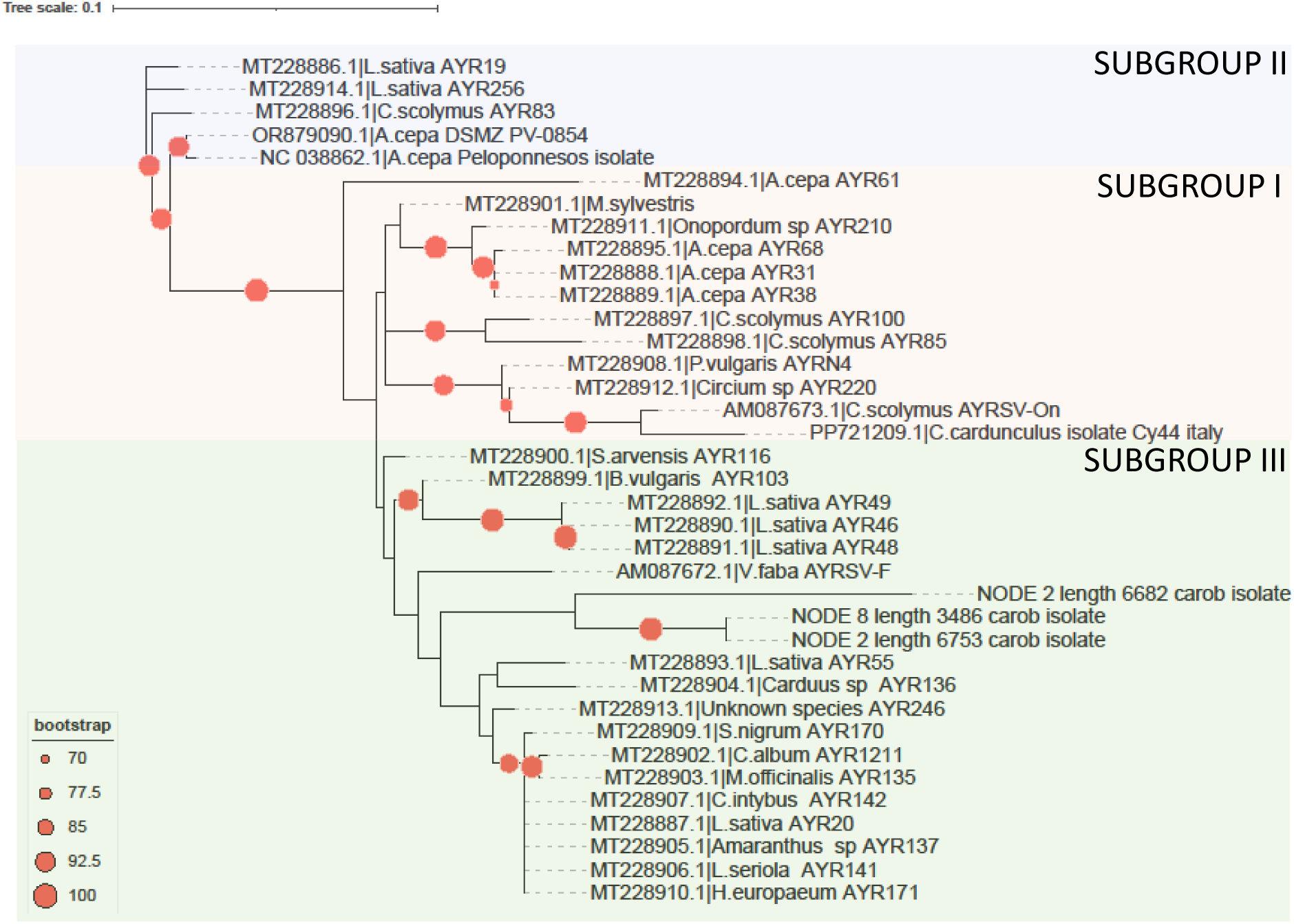
Unrooted phylogenetic analysis of AYRSV isolates using the Maximum-Likelihood algorithm (1000 bootstraps, excluding values below 75%). The region used is a partial RdRp region that includes motif A and an upstream region, according to Karapetsi *et al*. [25].

#### Strawberry latent ringspot virus

Strawberry latent ringspot virus (SLRSV) (species *Stralarivirus fragariae*) is part of the relatively recent Stralarivirus genus, together with Cohombrillo-associated virus (species *Stralarivirus elaterii*), Beldersay stralarivirus 1 (species *Stralarivirus beldersayense),* Lychnis mottle virus (species *Stralarivirus lychnis*) and beach cabbage stralarivirus (species *Stralarivirus scaevolae*). SLRSV was identified in 1964 and its host range exceeds 11 families, including economically important crops like fruits (strawberry, raspberry, blackberry, peach, plum), vegetables (celery, asparagus, parsley), ornamentals (lily, rose), grapevine, and olive [37, 38]. However, a 1969 article suggests that the virus also infects trees like Aesculus, Euonymus, and Robinia spp [39]. SLRV is a non-enveloped virus of 30 nm, comprising a RNA1 of 7.5Kb and a RNA2 of 3.8Kb, both with a VPg at the 5’ end and a polyA tail at the 3’end. This virus is transmitted by the soil-inhabiting nematodes *Xiphinema coxi* and *X. diversicaudatum,* however, seed transmission has also been proposed [38, 40]. SLRSV is mostly found in Europe, but in the last few years, incidences have been detected in the United States, Canada, North Africa, Taiwan, and New Zealand (https://gd.eppo.int/taxon/SLRSV0/distribution). Usually, this virus produces no symptoms, but yellow vein banding was observed in *Mentha sp.*, chlorotic spots on *Anemone sp*. and *Rubus spp.*, chlorotic streaks and necrotic rings on *Impatiens walleriana*, chlorotic mottle in *Solanum myricatum,* and finally vein chlorosis on Tibouchina sp. [38, 41]. An increase in the observed symptoms has also been described in the case of mixed infection of SLRSV with strawberry pallidosis-associated virus (species *Crinivirus palidofragariae*) and beet pseudo-yellows virus (species *Crinivirus pseudobetae*) [42]. In Greece, this virus has been identified only in Olive trees, including trees from the Chania region [43].

In this study, we identified 250 contigs across almost all pools tested, ranging from 200 nt to 7,218 nt, showing 65.83 % –91.23 % similarity to known isolates of the *Stralarivirus* family (primarily SLRSV), indicating a widespread presence of the virus in carob trees in Crete (Table S6). These contigs corresponded to both RNA1 and RNA2 of the viral genome. SLRSV is a virus presenting high nucleotide-level variability, which increases the difficulty in verifying the presence of the virus in each individual sample. To address this variability, we designed several primer sets, all presented in Table S3, and performed multiple different PCR amplifications to secure the presence of the virus (Figure S5). Most of the PCR products were Sanger sequenced, and fragments presented a similarity between 98.21 % and 100 % with the obtained contigs (Table S7). In total, we identified 53/134 positive samples (detection frequency of 39.55 %), with most of the samples located in Rethymnon (25), followed b y 15 in Heraklion, 9 in Chania and only 4 in the area of Lasithi (Table S5). This distribution indicates that the virus is detected more frequently in Rethymnon, suggesting that the area may represent a focal point of its spread.

Regarding the observed phenotypes, all collected tissues were examined, and only minimal leaf yellowing was detected, mainly around the veins. However, this symptom was subtle and not a reliable or easily distinguishable phenotype (Figure S6).

According to Dullemans *et al.*, SLRV isolates cluster into at least three distinct phylogenetic clades, named SLRSV-A, SLRSV-B and SLRSV-C [44]. Therefore, we examined the placement of our isolates within these established clades through a phylogenetic analysis. According to the ICTV, both the PRO-POL protein and the CP protein need to be assessed for a phylogenetic analysis. We repeated the analysis by Dullemans and included newly identified isolates from NCBI as well as our isolates. Figure 5 shows two main findings. First, we identified a new cluster (termed SLRSV-D) that groups with sequences from Dullemans work as well as newly submitted sequences. This difference may reflect the use of a different phylogenetic algorithm in our analysis (maximum likelihood rather than neighbor-joining) as well as the inclusion of additional SLRSV sequences. Second, our carob SLRSV isolates do not cluster within any of these four clades, in both the PRO-POL (region between CG and GDD) and CP (large and small) proteins. Instead, they form one additional, well-supported clade, which we propose as SLRSV-E. Within this clade, two well-distinguished subclades are observed, providing a clear explanation for the challenges encountered in designing uniform primers that amplify all samples. A consistent phylogenetic pattern was obtained when the sequences were analyzed at the nucleotide level (Figure S7). Taken together, these results suggest that genetically distinct SLRSV isolates were detected in carob, a previously unreported host.

**Figure 5:**
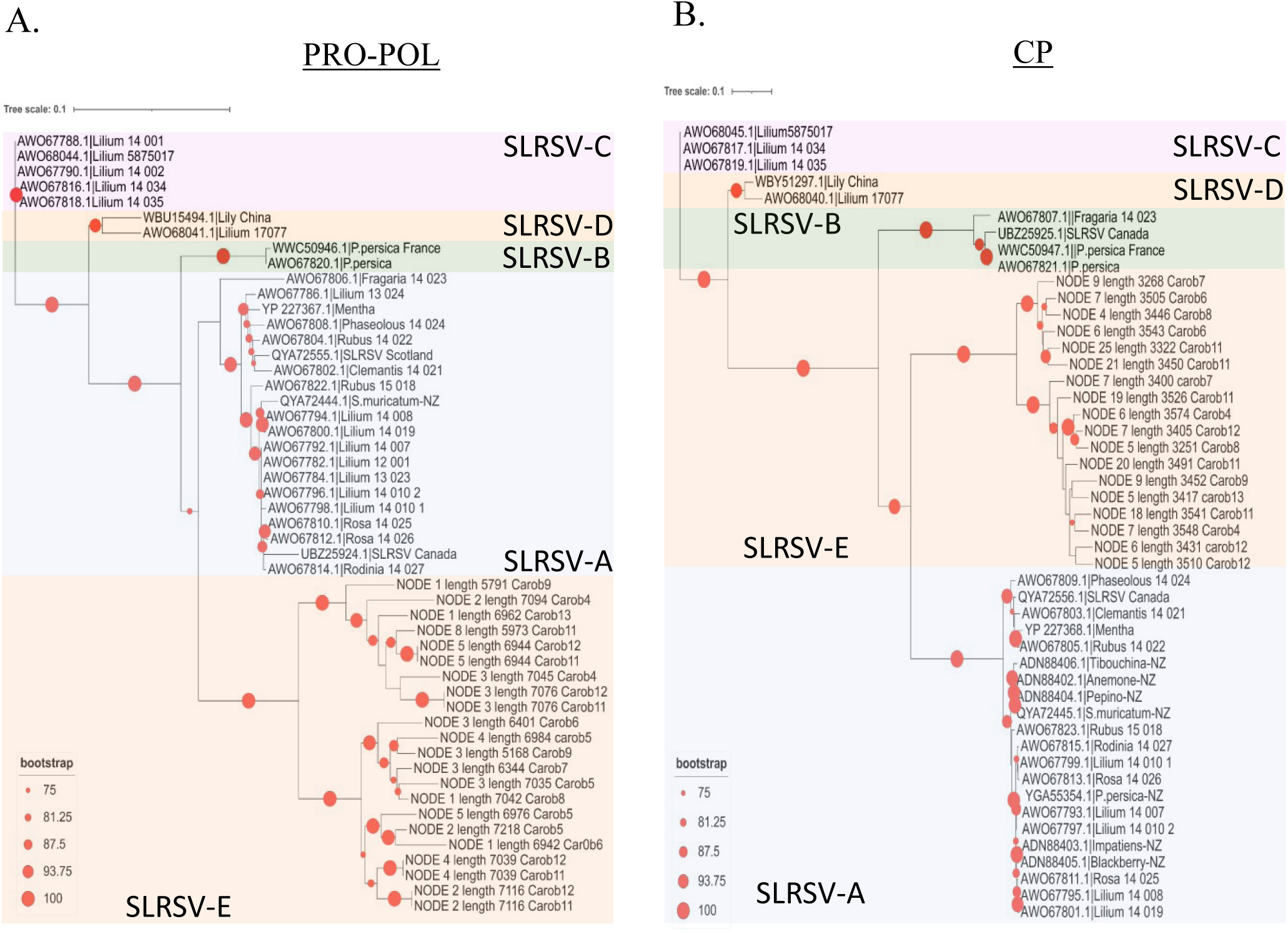
Unrooted intraspecies phylogenetic analysis of SLRSV isolates using Maximim-likelihood algorithm (1000 bootstraps, excluding values below 75%) for A) the ProPol region between the CG motif of the proteinase and the GDD motif of the polymerase and B) CPs, both large and small, of the virus according to Dullemans *et al.*, [44].

### Family *Bromoviridae*

*Bromoviridae* is a large family of tripartite, positive-sense RNA viruses enclosed in spherical, semispherical or bacilliform particles [45]. The family contains six genera, including *Ilarvirus*, a group first identified in the early 1900s [46]. The family name was originally created for viruses producing isometric, <u>la</u>bile particles associated with ringspot symptoms, and was later adopted by the ICTV [46]. Ilarviruses infect mostly woody plants such as prunus species, with Prunus necrotic ringspot virus (species *Ilarvirus PNRSV*), Prune dwarf virus (species *Ilarvirus PDV*), Apple mosaic virus (species *Ilarvirus* ApMV), and American plum line pattern virus (species *Ilarvirus APLPV*) being the best-known representatives [47]. Nevertheless, a wide range of other fruit crops and ornamentals have also been reported to host various ilarviruses [48, 49]. Symptoms range from latent infection to leaf mottling, deformation, stunting, fruit disorders, and even plant decline in some hosts [47, 49, 50]. Ilarviruses are transmitted by pollen, seeds and propagation material making its control especially challenging [47].

In this work, we identified 9 contigs (230 - 1265 nt) in one of the tested pools that showed resemblance to Soybean ilarvirus 1 (species *Ilarvirus SolV1*, also referred to as SIlV1 in the original report) with nucleotide similarity ranging from 86.8 % to 98.1 % (Table S8). This virus has three RNAs of around 3.6 Kb, 2.9 Kb and 2.3 Kb respectively [51]. RNA1 codes for the methyltransferase and the helicase of the virus, RNA2 for the RdRp and two more open reading frames (ORFs) of unknown function, and RNA3 for the movement and coat proteins [51]. Contigs corresponding to all three genomic RNA segments were recovered, further supporting the presence of this virus in carob trees.

We used specific primers (Table S3) for PCR and identified the M2 tree in the Lasithi prefecture containing the virus, as shown in Figure 6A. The PCR product was Sanger sequenced, and an identity of 100 % with its respective contigs was observed. No obvious foliar symptoms were observed in the SolV1-positive M2 tree (Figure 6B). SolV1 was first identified in the USA, but its geographical distribution remains unclear [51].

**Figure 6:**
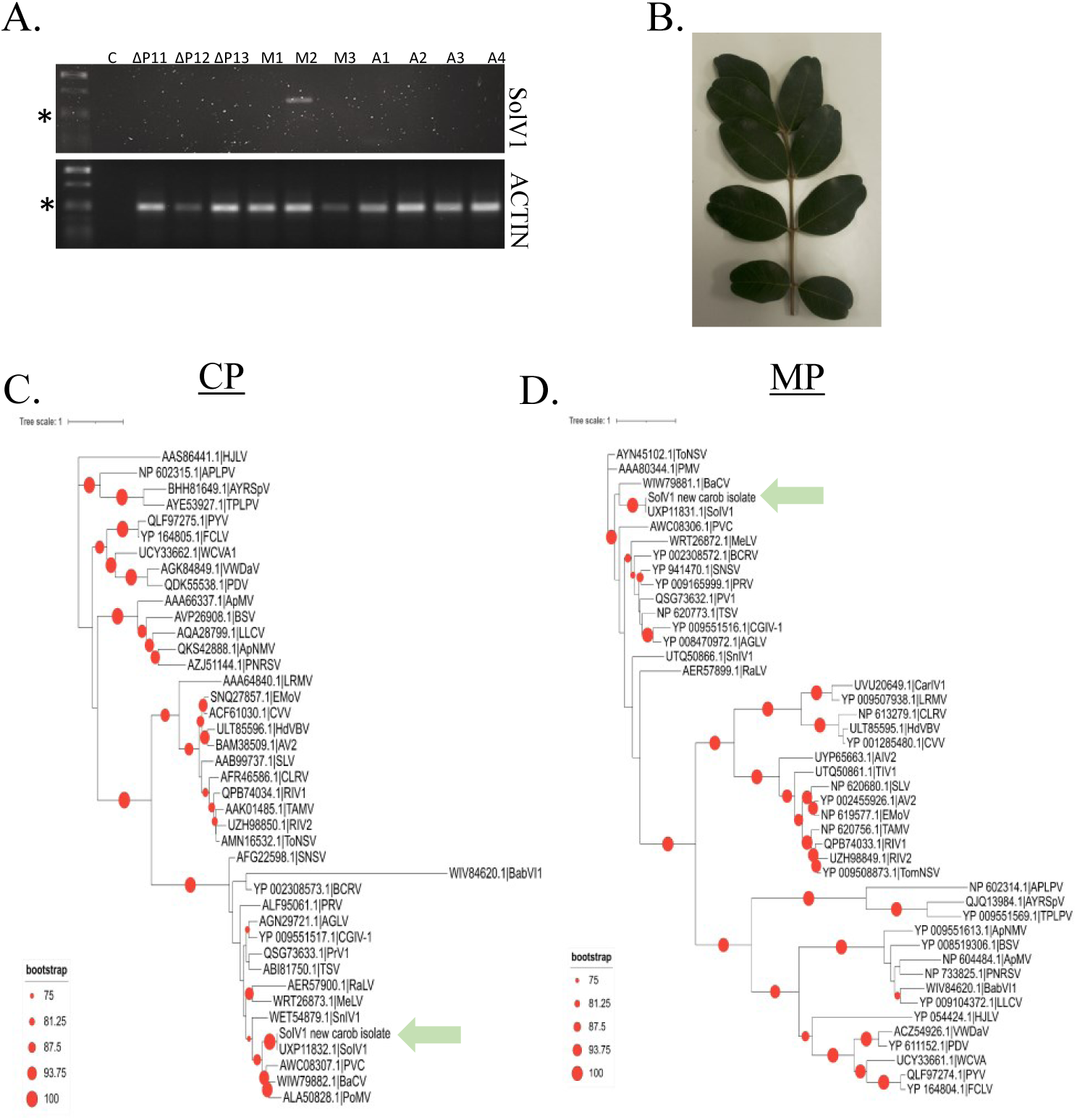
Identification of SolV1 in carob trees. A) PCR results from the individual samples belonging to the pools in which specific contigs were identified. Actin was used as an internal control to assess sample quality. The (*) corresponds to the 514bp band of λ phage/*Pst*I marker. ‘c’ stands for ‘control’ indicating the presence of water and not template during the PCR. B) Phenotypes observed in the SolV1-infected M2 plant. C and D) Unrooted intragenus phylogram for the CP and the MP proteins of various Ilarviruses according to Elmore *et al.*, [51].

It has been proposed that Ilarviruses can be divided into 4 subgroups, based on serology, sequence similarity and biological traits [46, 52]. As noted above, contigs corresponding to all three genomic RNA segments were detected; however, near-complete sequence coverage was recovered only for the CP and MP coding regions of RNA3. Therefore, we performed phylogenetic analysis using the sequences examined in Elmore *et al.*, together with newly identified sequences from recently described ilarviruses, using the CP and the MP of our isolate. As shown in Figure 6C and D, our isolate, as expected, is clustering with the only available isolate of SolV1 identified in the USA. It is also very closely linked to the viruses peanut virus C, bacopa chlorosis virus and potato mottle virus identified in the Republic of Korea, Germany and Spain respectively. To our knowledge, this is the first report of SolV1 in Greece and Europe, as well as the first detection of this virus in carob.

### Putative jivivirus

Jiviviruses are a group of single-stranded positive multipartite RNA viruses. These viruses have affinities with the insect-infecting Jingmen-like viruses (resembling *Flaviviridae*) and the plant-infecting virgaviruses [53]. This group has not yet been formally recognized by the ICTV committee, since only a few members have been described till today. The first jivivirus was identified in 2017 in citrus trees exhibiting citrus sudden death symptoms; however, the authors were unable to link the *Flaviviridae* and *Virgaviridae* contigs to a single infectious agent [54]. The proposal for this new group of viruses originated from the analysis of grapevine samples infected with *Plasmopara viticola*, in which conserved regions were detected at the 5′ and 3′ ends of several contigs. The newly identified virus was named grapevine-associated jivivirus 1 (GaJV-1), and the authors introduced the term “jiviviruses” by combining *Jingmen* and *Virga* (JiVi) [55]. Since then, more Jiviviruses have been identified in grapes, citrus and carya [53, 56].

During the virome analysis of carob trees, we identified 4 contigs from the same pooled sample resembling known Jiviviruses (e.g., pecan-associated Jivivirus 1), showing nucleotides similarity from 70.34 % to 87.93 % (Table S9). To confirm the presence of the virus, we performed PCR using specific primers and detected the virus only in sample T2 (Table S3-Figure S8). Sanger sequencing of two amplicons showed 99.6 % and 99.51 % identity, respectively, confirming that the virus is indeed present in this tree. We also identified fragments from various virgaviruses and Jingmen-like viruses. Despite these results, we were unable to recover the full-length viral genomes, which limits our ability to characterize this putative new virus or infer its biological properties. When examining the trees’ phenotype, we did not observe any notable symptoms apart from galls on the main branches, which, however, can be produced by numerous other factors apart from the presence of this virus (Figure S8). Taken together, our findings indicate the presence of a putative jivivirus in a sampled carob tree, although additional sequencing and biological analyses are required to determine its complete genome, evolutionary placement, and potential impact on tree health.

### Family *Phenuiviridae*

*Phenuiviridae* contains multipartite (2 to 8 segments) negative-sense or ambisense RNA viruses with a total genome size of up to 25 kb. Most members are enveloped, with non-enveloped members typically infecting plants, fungi, as well as arthropod vectors [57]. According to ICTV till today, there are 23 genera in this family, one of which is Coguvirus containing 8 viruses: Brassica campestris chinensis coguvirus 1 - BCCoV1 (species *Coguvirus chinense*), Edgeworthia chrysantha mosaic-associated virus - ECMaV (species *Coguvirus chrysanthae*), citrus concave gum-associated virus - CCGaV (species *Coguvirus citri*), watermelon crinkle leaf-associated virus 1 - WCLaV-1 (species *Coguvirus citrulli*), citrus virus A (CiVA) (species *Coguvirus eburi*), watermelon crinkle leaf-associated virus 2 - WCLaV-2 (species *Coguvirus henanense*), blackberry line pattern virus - BlaLPV (species *Coguvirus rubi*) and Yúnnán Paris negative-stranded virus –YPNSV (species *Coguvirus yunnanense*). Two more viruses have been recently proposed (Brassica oleracea torzella virus 1 –BoTV1 (species *Coguvirus torzellae*), and yellow silver pine associated phenui-like virus –YSPaPLV) but are still unclassified [58]. Currently recognized hosts of this genus belong to the families *Brassicaceae*, *Cucurbitaceae*, *Melanthiaceae*, *Vitaceae*, and *Rutaceae* [58, 59]. Symptoms can range from mild (asymptomatic plants) to severe, with symptoms like mosaic, chlorosis, yellow spots, leaf shrinkage, fruit rind symptoms, and possibly citrus impetratura [59–63]. In Greece, CiVA, WCLaV-1 and WCLaV-2 have been found infecting citrus trees and watermelons, respectively [60, 61]. To date, only seed and graft transmission have been demonstrated, although additional transmission routes cannot be excluded. The genome of coguviruses consists of two RNAs. RNA1, typically ∼6.8–7.2 Kb in length, encodes RdRp. RNA2, usually ∼2.7–3.1 kb, is ambisense and encodes the movement protein and the capsid protein, separated by a long AU-rich intergenic region, a characteristic molecular feature of this genus.

In the HTS data, we identified 87 phenuivirus and coguvirus-like sequences ranging from 201 nt to 6888 nt (Table S10). BLASTn analysis showed low overall similarity to known viruses (approximately 68 %), particularly for the larger fragments, most of which aligned more closely with BCCoV1 [64]. When BLASTx analysis was used for the proteins produced from the largest contigs, it revealed low identity: 56.56 % for the RdRp protein (98 % coverage), 43.07 % for the movement protein (MP; 97 % coverage), and 59.18 % for the nucleocapsid protein (NP; 72 % coverage). According to ICTV criteria, an RdRp identity below 95% suggests the existence of a novel virus.

Because of the large number of contigs and the presence in almost all pools, we reasoned that the virus must be frequently detected in carob trees. We therefore used the primer pair CogD-F/CogD-R to screen all 134 collected samples for the presence of the virus (Table S3). As shown in Figure S9, 47 out of the 134 samples were identified as infected with this coguvirus, meaning that around 35 % of the tested plants were infected with this new virus in all four prefectures.

The symptoms associated with coguvirus infection have not been fully characterized for all members of the genus and range from absent or mild to highly pronounced [58, 60–63]. In this study, we identified several carob trees containing a new coguvirus and investigated a possible phenotype. As shown in Figure S10, no obvious phenotype was observed in the leaves of the studied carob trees, suggesting that this virus may be latent in this host, however, being able to exclude a possible effect in carob tree production.

Next, we designed a series of overlapping primers to amplify the complete genome from sample ΔΡ5 (Table S3-Figure S11). Although we attempted both 3′ and 5′ RACE, these efforts were unsuccessful. Therefore, we relied on contigs obtained from all samples and identified the probable terminal regions consistent with those previously described for members of the genus *Coguvirus*. Based on this analysis, RNA1 was determined to be 6730 nt in length and encodes a protein of 2213 aa (∼255 KDa), which is slightly larger than the L proteins reported for other coguviruses, as noted by ICTV (Figure 7A). The L protein shows the highest similarity to CCGaV (56.56 %, 98 % coverage), however, the relatively low identity leaves open the possibility that this virus may not belong to the coguvirus group. To address this, we examined whether the protein contains characteristic features of coguvirus L proteins. First, the L protein includes a ‘Bunya-RdRp domain’(positions 563-1235 aa), consistent with the domain architecture described for other members of the family [59, 65]. Second, five conserved motifs (A-E) have been proposed for the major representatives of this genus, and we therefore assessed their presence in our newly identified virus [65]. Figure 7B shows that the L protein contains all five proposed conserved motifs. For instance, motif C includes the essential SDD residues (position 1075 aa), typical of various negative-sense RNA (nsRNA) viruses including coguviruses [58, 59, 65, 66]. In addition, L protein contains a putative cap-snatching motif, a putative endonuclease domain (H_66_, D_71_, PD_95−96_, ExG_107_, K_126_) (Figure S12), found in many viruses of the *Orthomyxoviridae* family [67]. Taken together, these observations suggest that the L protein identified in this new virus likely functions as an RdRp and despite its relatively low similarity to known viruses, it retains all characteristic features of this genus.

**Figure 7:**
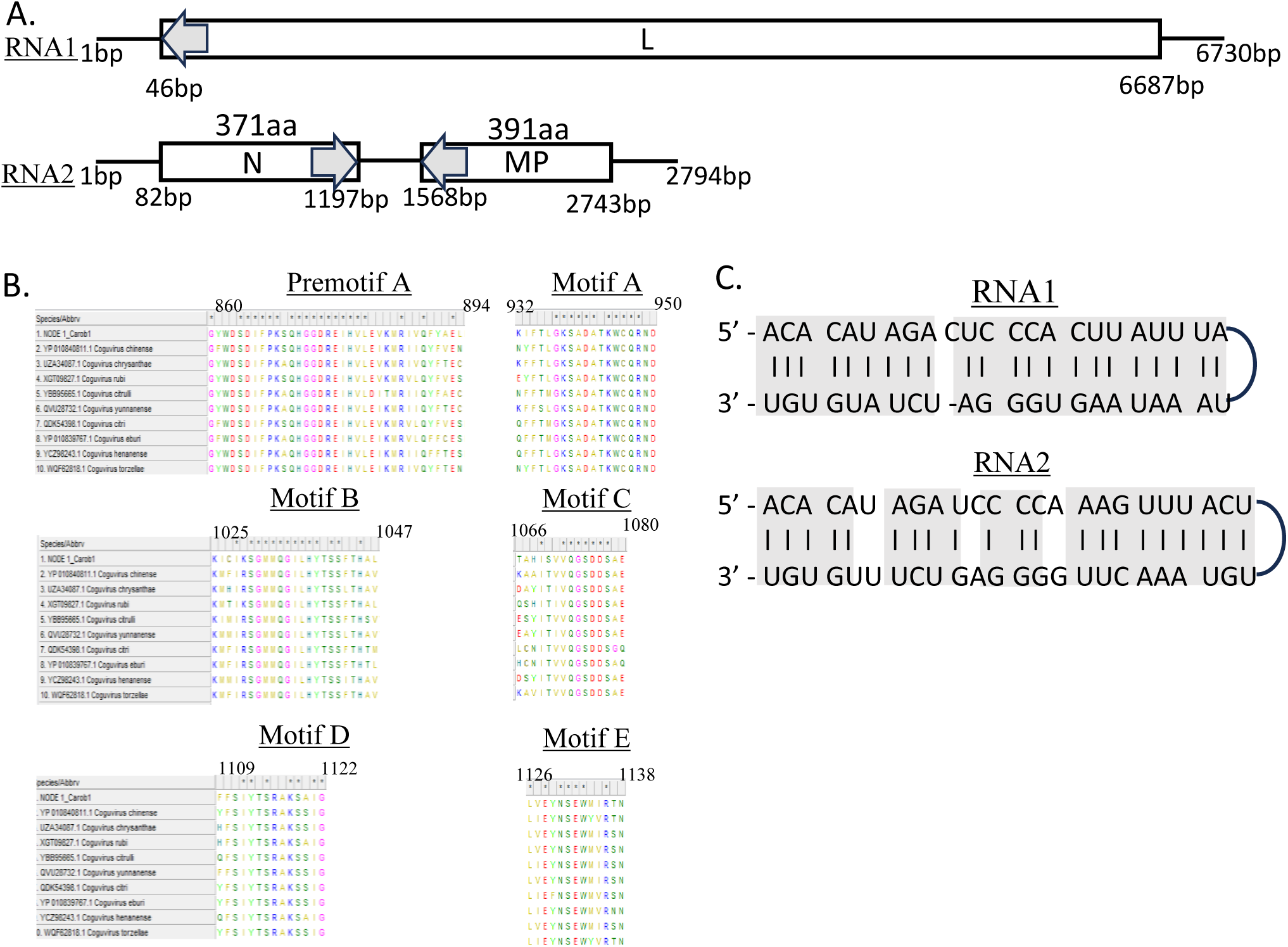
Novel coguvirus. A) Schematic representation of the new coguvirus (provisionally named Carob coguvirus 1 –CaCoV1). B) Sequence of the five motifs identified in major coguviruses according to [58, 65]. The numbering was performed using the aa of the L protein of the newly identified coguvirus. C) Panhandles created at the viral RNAs termini of CaCoV1.

The RNA 2 is 2794 nt long, and exhibits an ambisense genomic organization, encoding for two proteins: a nucleocapsid protein (N) produced from a 1176 nt ORF, corresponding to 371 aa (∼40 KDa), and a movement protein (MP), produced by a 1116 nt ORF, corresponding to 391 aa (∼44 KDa). These areas are separated by a 371 nt AU-rich intergenic region (approximately 70 % AU content), which, according to RNA folding predictions, could result in a highly structured region, as has been proposed for other coguviruses. The N protein has higher similarity with the WCLaV-1 N protein (59.18 %, 72 % coverage), which, as in the case of the L protein, is a relatively low similarity. N protein contains a region (134-302 aa) resembling Tenuivirus/Phlebovirus nucleocapsid protein, a characteristic found in other coguviruses.

The MP protein also shows low similarity to the CCGaV MP protein (43.7 %, 97 % coverage), and no known functional domain could be identified in its sequence. This further supports the notion that the virus represents a distinct member of the Coguvirus genus. Although no studies have described conserved domains specific to coguvirus movement proteins, alignment of MP sequences from a broad set of known coguviruses revealed that all of them, including MP of our newly identified virus, share conserved amino acids at specific positions within the central region of the protein, whereas the N-terminal region remains highly variable (Figure S13). Specifically, three short sequence motifs GxxFQYxA (position 72-79), LxDxR (position 163-166), and NSNxxxKxELS (position 182-192), were consistently observed across multiple coguvirus MPs. Even though no functional role has been assigned to those motifs in viral or cellular proteins according to the literature, it has been shown that the ‘Y’ of the first domain, the ‘D’ of the second motif and the ‘K’ of the third domain are conserved not only in coguvirus but also in other nsRNA viruses [65]. Nevertheless, additional investigation will be required to elucidate whether these conserved sequence features contribute to the function of these movement proteins.

The final aspect that we investigated for this new virus was the structure of its RNA ends. Previous studies have proposed that both genomic RNAs of coguviruses contain complementary sequences at their respective 5′ and 3′ ends, forming panhandle structures [58, 59, 65]. By analyzing all obtained contigs, we identified conserved 21 nt and 24 nt regions at the 5 ′ end of the two RNAs that are complementary to the sequences at the 3 ′ end of the same RNA, generating panhandle structures (Figure 7C). Moreover, these terminal sequences are highly similar to those reported for other members of this family (Figure S14).

We also performed phylogenetic analyses of the RdRp, N, and MP proteins of this new virus and compared them with those of known viruses in this genus. As shown in Figure 8, the new coguvirus is clearly found to be different but closely related to BoTV1 (species *Coguvirus torzellae*) from Italy and BCCoV1 (species *Coguvirus chinense*) from China for both the full region of RdRp and N proteins. In contrast, the MP of this new virus seems more closely related to members of Bociviruses, a different genus in this family than coguviruses, suggesting its structural resemblance to proteins from the tobacco mosaic virus 30 kDa family [68]. To further support that the identified virus is indeed new, we used SDT analysis. As shown in Figure S15 and Table S11, all proteins were verified with low similarity percentages to other proteins.

**Figure 8:**
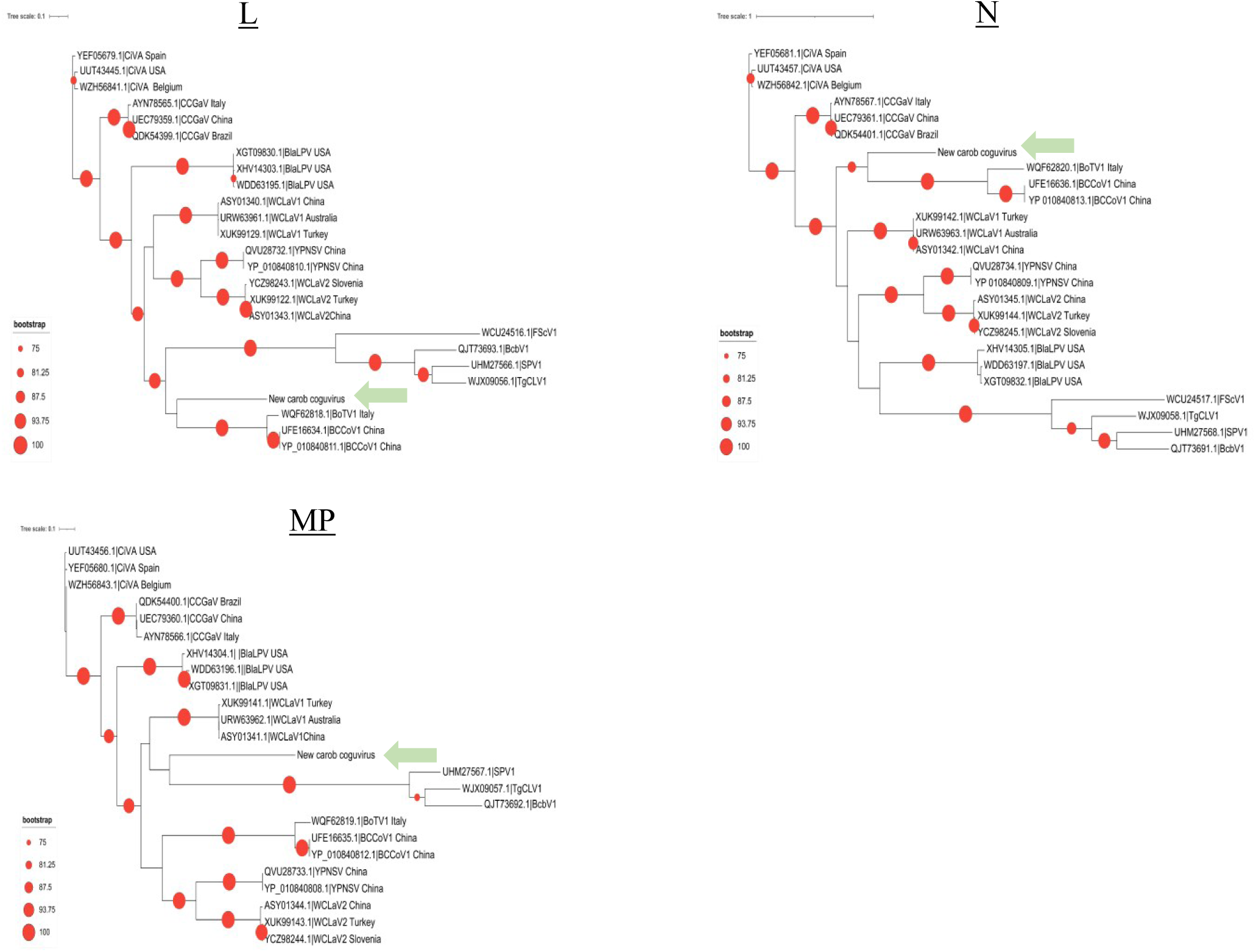
Phylogenetic analysis of various Coguviruses and Bociviruses proteins, specifically, L (A), N (B) and MP (C), using Maximum-likelihood algorithm with 1000 bootstraps and excluding values below 75%.

Taken together the analysis described above supports the presence of a new virus infecting carob trees that belongs to the genus Coguvirus within the *Phenuiviridae* family. We propose the provisional name Carob coguvirus 1 (CaCoV1, species *Coguvirus ceratoniae*).

## CONCLUSION

Carob is gaining renewed importance as a Mediterranean crop, particularly in the context of agricultural policies promoting more sustainable water use. By working directly with growers in Crete and examining both cultivation practices and tree health, we were able to build the first clear picture of the carob virome in the region. Our analyses revealed genetically distinct isolates of AYRSV and SLRSV, the first detection of Soybean ilarvirus 1 (SolV1) in Greece, a putative jivivirus with an incomplete genome, and a previously undescribed coguvirus, for which we propose the name Carob coguvirus 1 (CaCoV1). These findings show that carob trees host a far richer viral diversity than previously appreciated. To our knowledge, this is the first comprehensive virome-focused study of carob trees. It provides a foundation for future epidemiological research, virus surveillance, and the development of improved management strategies for this increasingly valuable Mediterranean crop.

## AUTHOR STATEMENTS

### AUTHOR CONTRIBUTIONS

KK, GE and KrK contributed to the conceptual development of this project and supervised the work. KK and GE developed the experimental strategy and collected the samples. KK, MB, AF organized the sample processing and performed experiments. GP and CA performed bioinformatic analyses. DM was responsible for participant follow up with participants and analyzed the obtained questionnaires. KK, MB, GE and KrK wrote the article and all authors critically revised the manuscript for scientific accuracy, clarity, and structure.

### CONFLICT OF INTEREST

The authors declare no competing interests.

### FUNDING INFORMATION

This work has been supported by a grant from the University of Crete (Category C) entitled ‘Phytopathogenic threats and economic viability of *Ceratonia siliqua* (Carob) production in Crete –[PhytoCARE] –KA11667 –Special account of Research Funds of University of Crete.

### ETHICAL APPROVAL

Written informed consent was obtained from all participants, and the study was approved by the Ethics Committee of the University of Crete (23/09.02.2024), according to the principles of the Declaration of Helsinki.

### DATA AVAILABILITY

Data from this work have been uploaded in Sequence Read Archive (SRA - NCBI) under the following ID: SUB16460380

## ACKNOWLEDGEMENTS

The authors would like to acknowledge all the participants who gave us their valuable time and material. We would also like to thank Korina Miliaraki and the Epimenidis Cultural Society in Panormo, Rethymnon (https://www.epimenides.gr) for their help in identifying carob growers. The authors would like to thank Giannis Pyrgianakis and Dr. Anastasia Kampouraki for helpful discussions concerning insects identified in the carob trees, and Lidia Nikolopoulou for technical assistance.

## Supplemental Figures

**Figure S1:** Denaturing agarose gels for the assessment of RNA quality obtained from the collected leaves.

**Figure S2:** Phenotypes observed in the collected carob samples. Based on the literature, these symptoms may be associated with fungal infections (A), bacterial infections (B), or insect damage (C).

**Figure S3:** AYRSV in carob trees. PCR results from the individual samples belonging to the pools in which AYRSV-specific contigs were identified. Actin was used as an internal control for the quality of the obtained samples. The actin bands correspond to the same gel shown in multiple figures, as the same set of samples were used. The (*) corresponds to the 514bp band of λ phage/*Pst*I marker. ‘c’ stands for ‘control’ indicating the presence of water and not template during the PCR.

**Figure S4:** Representative pictures from leaves of AYRSV-infected carob trees identified in this work.

**Figure S5:** SLRSV in carob trees. PCR results from the individual samples belonging to the pools in which SLRSV-specific contigs were identified. Actin was used as an internal control for the quality of the obtained samples. The actin bands correspond to the same gel shown in previous figures, as the same set of samples was used. The (*) corresponds to the 514bp band of λ phage/*Pst*I marker. ‘c’ stands for the negative control including only water during the PCR.

**Figure S6:** Representative pictures from leaves of SLRSV-infected carob trees identified in this work.

**Figure S7:** Unrooted phylogenetic analysis at a nucleotide level (cds) for the regions of the ProPol (A) and the CP (B) for SLRSV viruses.

**Figure S8:** Possible new Jivivirus in carob trees. A) PCR results from the individual samples belonging to the pools in which Jivi-specific contigs were identified. T2 carob tree was found to contain this virus. Actin was used as an internal control for the quality of the obtained samples. The actin bands correspond to the same gel shown in previous figures, as the same set of samples was used. The (*) corresponds to the 514bp band of λ phage/*Pst*I marker. ‘c’ stands for ‘control’ indicating the presence of water and not template during the PCR. B) Representative pictures from carob tree T2.

**Figure S9:** Identification of the new coguvirus CaCoV1 in carob trees. PCR results from the individual samples belonging to the pools in which virus-specific contigs were identified. Actin was used as an internal control for the quality of the obtained samples. The actin bands correspond to the same gel shown in previous figures, as the same set of samples was used. The (*) corresponds to the 514bp band of λ phage/*Pst*I marker. ‘c’ stands for ‘negative control’ using only water during the PCR.

**Figure S10:** Phenotype in leaves from CaCoV1-infected carob trees.

**Figure S11:** Overlapping PCRs to sequence the full genome of the newly identified coguvirus, CaCoV1. The first well contains a λ phage/*Pst*I marker, the second a negative control using water instead of a template, and the third corresponds to the result of the studied PCR.

**Figure S12:** Cap-snatching motif identified in the L protein of various coguvirus according to the domain found in many *Orthomyxoviridae* [67]. The numbering follows the aa of the L protein of the newly identified coguvirus.

**Figure S13:** Motifs identified in the MP of various coguviruses. The numbering follows the aa of the MP protein of the newly identified coguvirus.

**Figure S14:** Sequence of the RNA termini of various coguviruses. By analyzing a large number of isolates (not shown here), we were able to show that our virus follows the ‘canonical’ sequences at the ends, creating panhandle sequences. For visualization, we used the weblogo software.

**Figure S15:** Pairwise sequence comparison of the L (A), N (B) and MP (C) proteins regions of the novel coguvirus, together with reference sequences of other published coguviruses and bociviruses (Table S11), using SDTv1.3 [29].

## Supplementary Tables

**Table S1:** Leaf collection dates

**Table S2:** HTS libraries reads

**Table S3:** Primers used in this study

**Table S4:** Contigs identified for AYRSV

**Table S5:** PCR verification results for each virus

**Table S6:** Contigs identified for SLRSV

**Table S7**: Sanger sequencing of SLRSV PCR products

**Table S8:** Contigs identified for SILV

**Table S9:** Contigs identified for a putative Jivivirus

**Table S10:** Contigs identified for a new Coguvirus

**Table S11:** SDTv1.3 pairwise analysis of the L, N and MP proteins of coguviruses and bociviruses assessed in this study

## Footnotes

1 All numbers correspond to 2017 production quantities and, except for Spain, are obtained from FAOSTA (https://www.fao.org/faostat/en/#data/QCL/visualize). Different vintages of the FAOSTAT database report very different values for Spain and, for this reason, the corresponding quantity is obtained from the statistics reported by the Spanish Ministry of Agriculture, Fisheries and Food (https://www.mapa.gob.es/estadistica/pags/anuario/2020/TABLAS%20PDF/CAPITULO07/pdfc07_7.13.2.1.pdf, Table 7.13.2.1).

2 The abrupt decline in the number of trees in the prefecture of Rethymno between 2013 and 2014 is most likely due to a change in the surveying and monitoring standards used by the Hellenic Statistical Authority.

## Notes

### Competing Interest Statement

The authors have declared no competing interest.

