## Supplemental Figures for "The Cretan Carob Virome": Supplementary Figures-Final.pdf

**Figure S1**

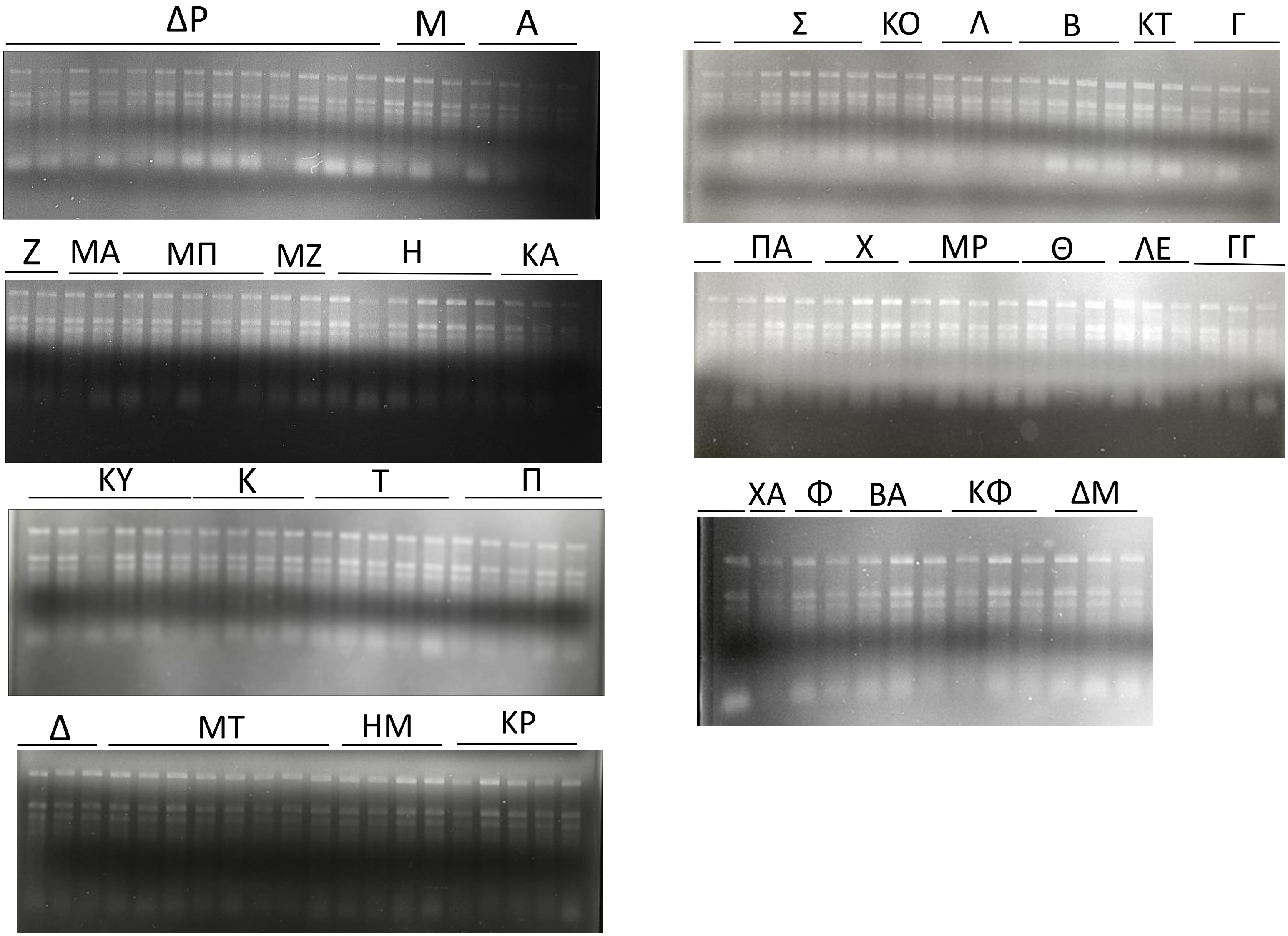
ΔΡ
Μ
Α
Σ
ΚΟ
Λ
Β
ΚΤ
Γ
Ζ
ΜΑ
ΜΠ
ΜΖ
Η
ΚΑ
ΠΑ
Χ
ΜΡ
Θ
ΛΕ
ΓΓ
ΚΥ
Κ
Τ
Π
ΧΑ
Φ
ΒΑ
ΚΦ
ΔΜ
Δ
ΜΤ
ΗΜ
ΚΡ

**Figure S2**

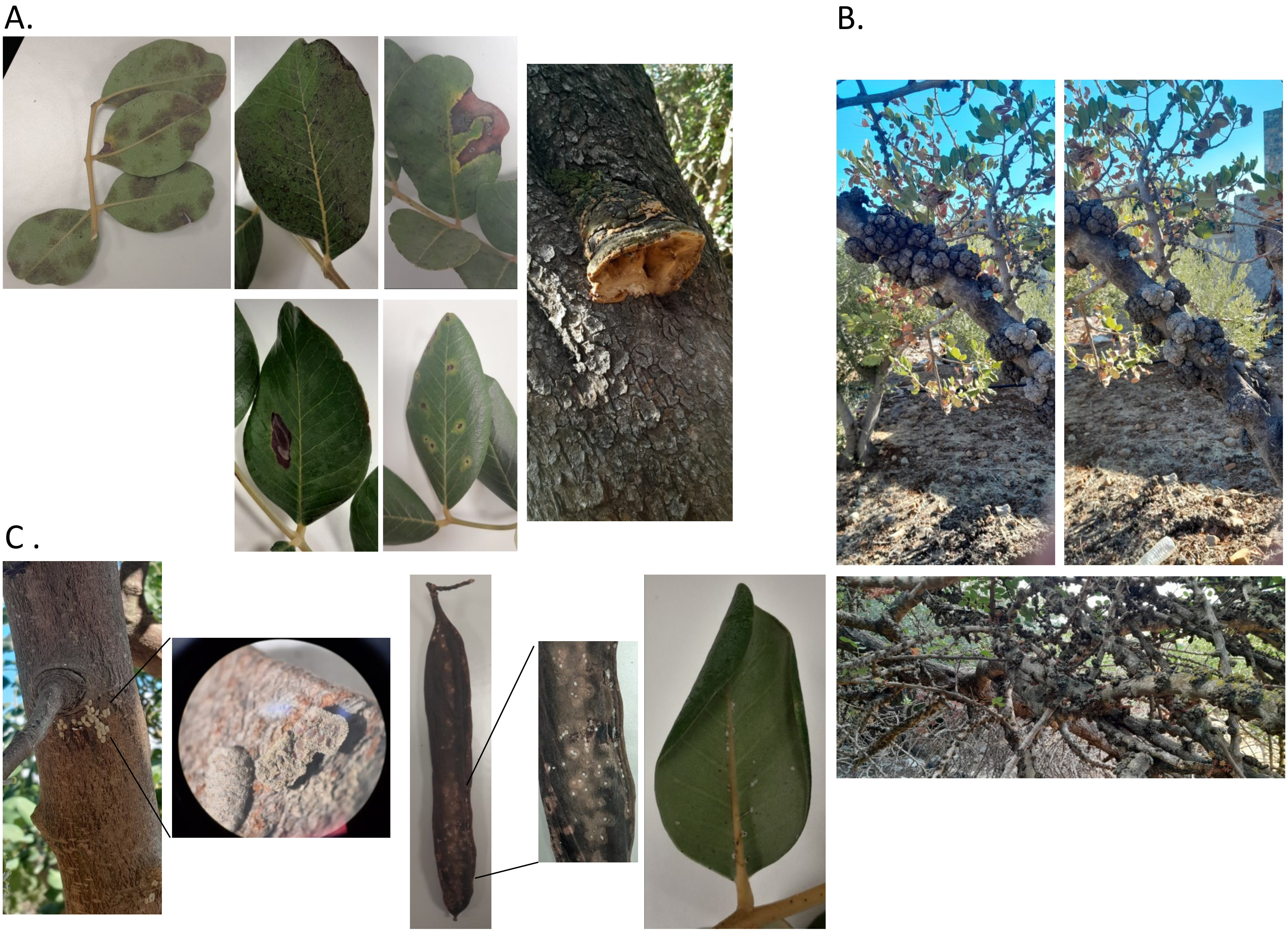

### Figure S3

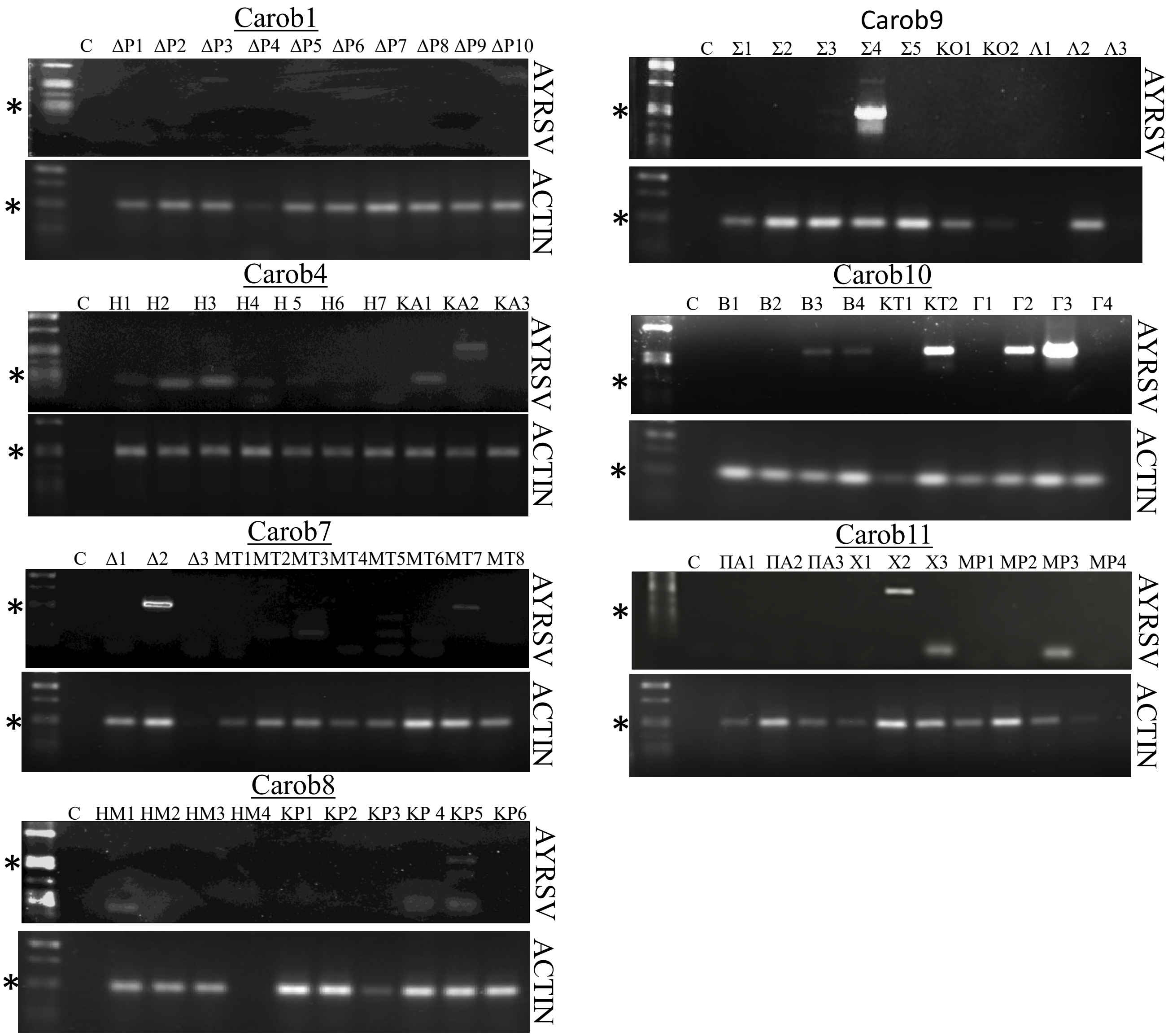

Carob1
C ΔP1 ΔP2 ΔP3 ΔP4 ΔP5 ΔP6 ΔP7 ΔP8 ΔP9 ΔP10
AYRSV
ACTIN
Carob4
C H1 H2 H3 H4 H 5 H6 H7 KA1 KA2 KA3
AYRSV
ACTIN
Carob7
C Δ1 Δ2 Δ3 MT1MT2MT3MT4MT5MT6MT7 MT8
AYRSV
ACTIN
Carob8
C HM1 HM2 HM3 HM4 KP1 KP2 KP3 KP 4 KP5 KP6
AYRSV
ACTIN
Carob9
C Σ1 Σ2 Σ3 Σ4 Σ5 KO1 KO2 Λ1 Λ2 Λ3
AYRSV
Carob10
C B1 B2 B3 B4 KT1 KT2 Γ1 Γ2 Γ3 Γ4
AYRSV
ACTIN
Carob11
C ΠA1 ΠA2 ΠA3 X1 X2 X3 MP1 MP2 MP3 MP4
AYRSV
ACTIN

**Figure S4**

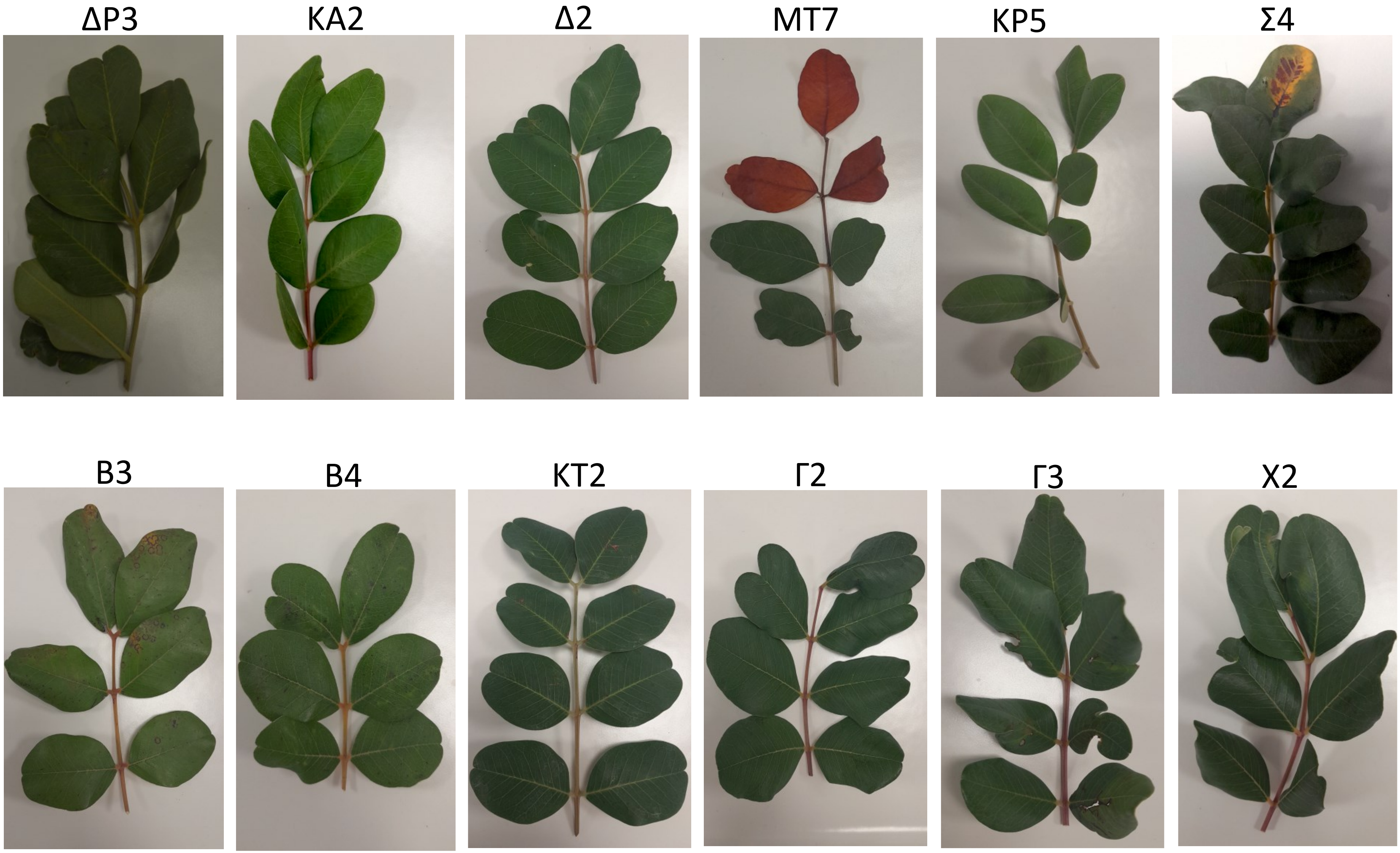

### Figure S5

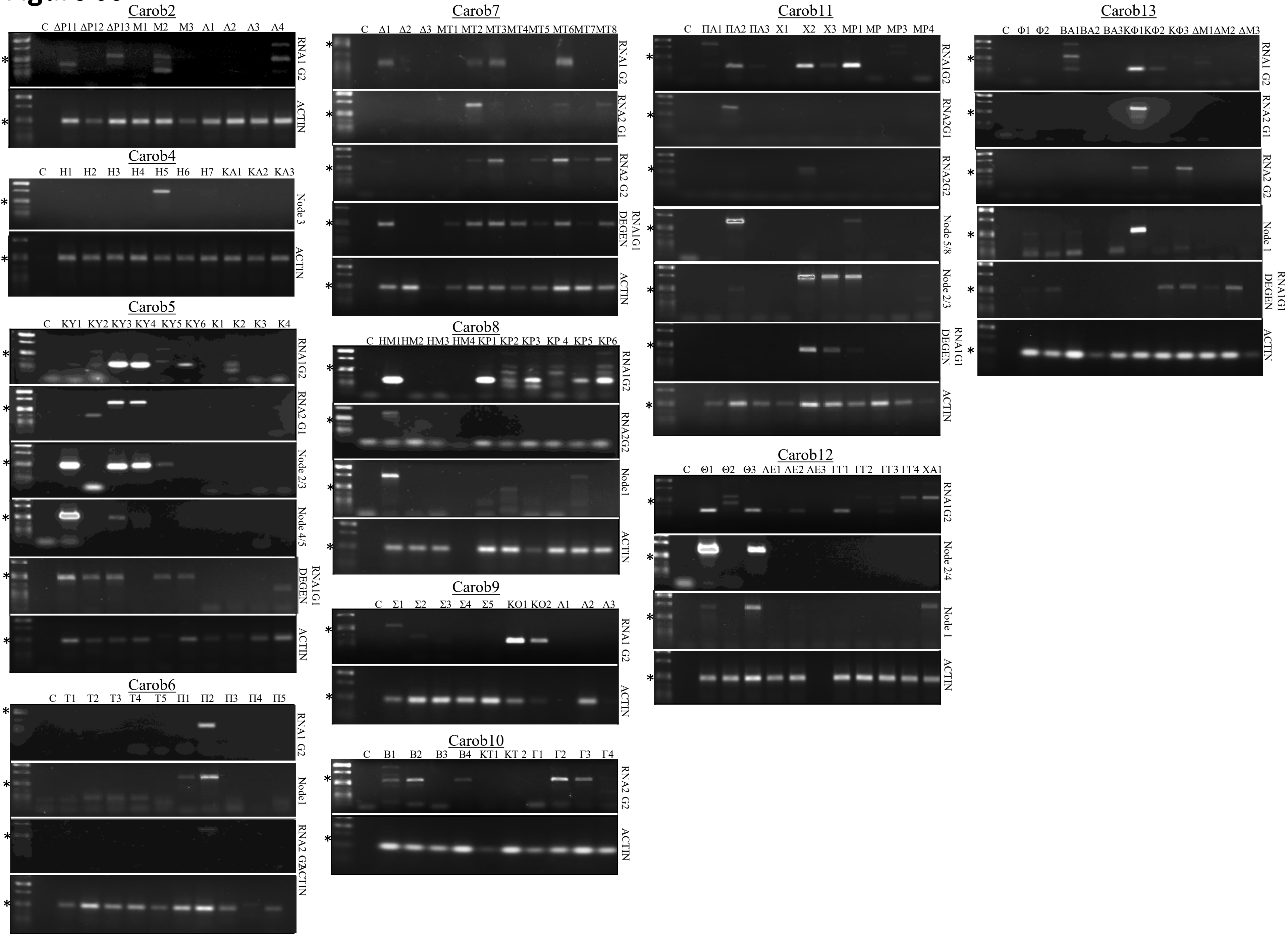
Carob2
C ΔP11 ΔP12 ΔP13 M1 M2 M3 A1 A2 A3 A4
RNA1 G2
ACTIN
Carob4
C H1 H2 H3 H4 H5 H6 H7 KA1 KA2 KA3
Node 3
ACTIN
Carob5
C KY1 KY2 KY3 KY4 KY5 KY6 K1 K2 K3 K4
RNA1G2
RNA2 G1
Node 2/3
Node 4/5
RNA1G1 DEGEN
ACTIN
Carob6
C T1 T2 T3 T4 T5 Π1 Π2 Π3 Π4 Π5
RNA1 G2
Node1
RNA2 G2
ACTIN
Carob7
C Δ1 Δ2 Δ3 MT1 MT2 MT3 MT4 MT5 MT6 MT7 MT8
RNA1 G2
RNA2 G1
RNA2 G2
RNA1G1 DEGEN
ACTIN
Carob8
C HM1 HM2 HM3 HM4 KP1 KP2 KP3 KP 4 KP5 KP6
RNA1G2
RNA2G2
Node1
ACTIN
Carob9
C Σ1 Σ2 Σ3 Σ4 Σ5 KO1 KO2 Λ1 Λ2 Λ3
RNA1 G2
ACTIN
Carob10
C B1 B2 B3 B4 KT1 KT 2 Γ1 Γ2 Γ3 Γ4
RNA2 G2
ACTIN
Carob11
C ΠA1 ΠA2 ΠA3 X1 X2 X3 MP1 MP MP3 MP4
RNA1G2
RNA2G1
RNA2G2
Node 5/8
Node 2/3
RNA1G1 DEGEN
ACTIN
Carob12
C Θ1 Θ2 Θ3 ΛE1 ΛE2 ΛE3 ΓΓ1 ΓΓ2 ΓΓ3 ΓΓ4 XA1
RNA1G2
Node 2/4
Node 1
ACTIN
Carob13
C Φ1 Φ2 BA1 BA2 BA3 KΦ1 KΦ2 KΦ3 ΔM1 ΔM2 ΔM3
RNA1 G2
RNA2 G1
RNA2 G2
Node 1
RNA1G1 DEGEN
ACTIN

**Figure S6**

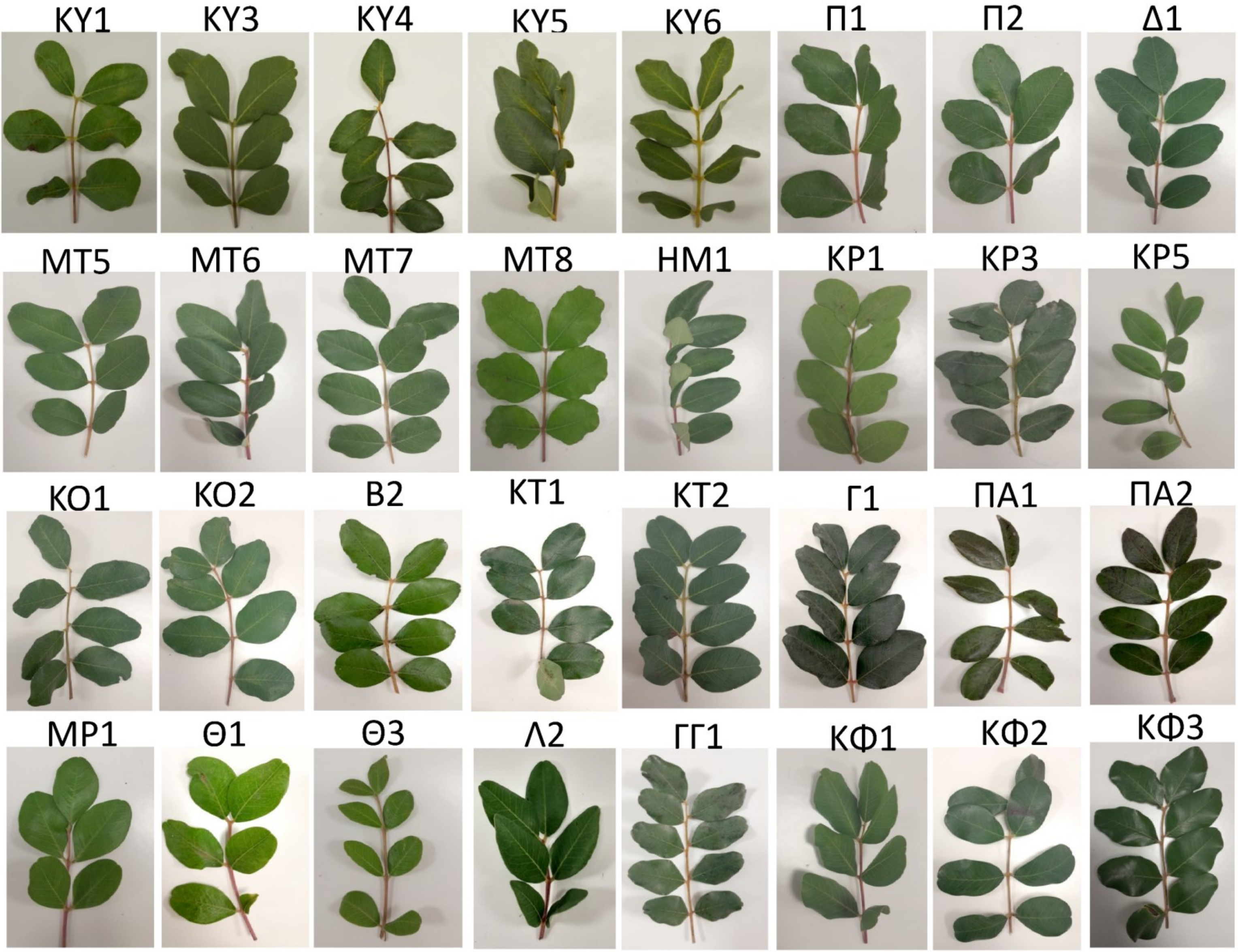
ΚΥ1
ΚΥ3
ΚΥ4
ΚΥ5
ΚΥ6
Π1
Π2
Δ1
ΜΤ2
ΜΤ3
ΜΤ5
ΜΤ6
ΜΤ7
ΜΤ8
ΗΜ1
ΚΡ1
ΚΡ3
ΚΡ5
ΚΡ6
Σ2
ΚΟ1
ΚΟ2
Β2
ΚΤ1
ΚΤ2
Γ1
ΠΑ1
ΠΑ2
Χ2
Χ3
ΜΡ1
Θ1
Θ3
Λ2
ΓΓ1
ΚΦ1
ΚΦ2
ΚΦ3

### Figure S7

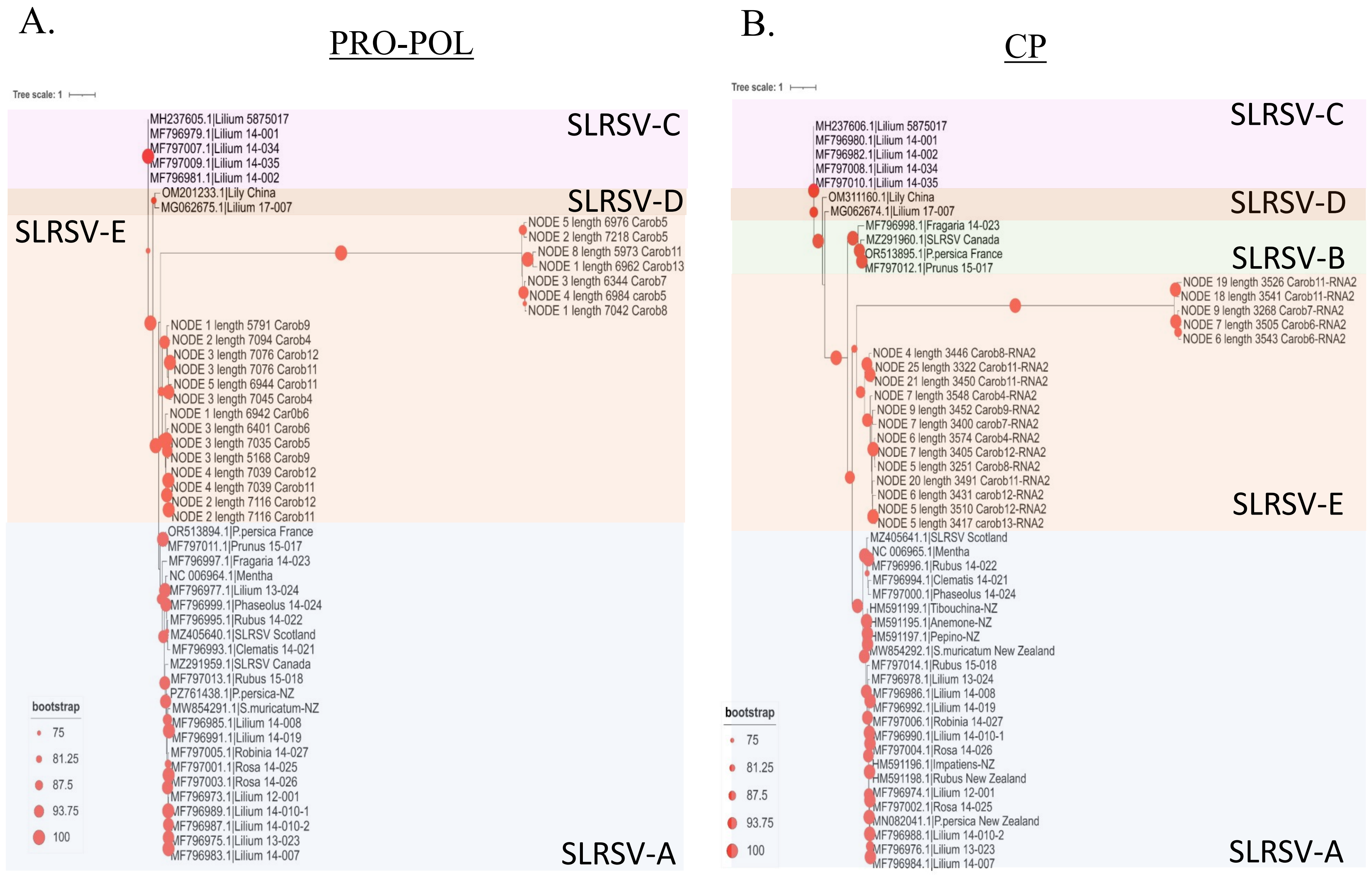
A.
PRO-POL
Tree scale: 1
SLRSV-C
MH237605.1|Lilium 5875017
MF796979.1|Lilium 14-001
MF797007.1|Lilium 14-034
MF797009.1|Lilium 14-035
MF796981.1|Lilium 14-002
SLRSV-D
OM201233.1|Lily China
MG062675.1|Lilium 17-007
SLRSV-E
NODE 5 length 6976 Carob5
NODE 2 length 7218 Carob5
NODE 8 length 5973 Carob11
NODE 1 length 6962 Carob13
NODE 3 length 6344 Carob7
NODE 4 length 6984 carob5
NODE 1 length 7042 Carob8
NODE 1 length 5791 Carob9
NODE 2 length 7094 Carob4
NODE 3 length 7076 Carob12
NODE 3 length 7076 Carob11
NODE 5 length 6944 Carob11
NODE 3 length 7045 Carob4
NODE 1 length 6942 Car0b6
NODE 3 length 6401 Carob6
NODE 3 length 7035 Carob5
NODE 3 length 5168 Carob9
NODE 4 length 7039 Carob12
NODE 4 length 7039 Carob11
NODE 2 length 7116 Carob12
NODE 2 length 7116 Carob11
OR513894.1|P.persica France
MF797011.1|Prunus 15-017
MF796997.1|Fragaria 14-023
NC 006964.1|Mentha
MF796977.1|Lilium 13-024
MF796999.1|Phaseolus 14-024
MF796995.1|Rubus 14-022
MZ405640.1|SLRSV Scotland
MF796993.1|Clematis 14-021
MZ291959.1|SLRSV Canada
MF797013.1|Rubus 15-018
PZ761438.1|P.persica-NZ
MW854291.1|S.muricatum-NZ
MF796985.1|Lilium 14-008
MF796991.1|Lilium 14-019
MF797005.1|Robinia 14-027
MF797001.1|Rosa 14-025
MF797003.1|Rosa 14-026
MF796973.1|Lilium 12-001
MF796989.1|Lilium 14-010-1
MF796987.1|Lilium 14-010-2
MF796975.1|Lilium 13-023
MF796983.1|Lilium 14-007
SLRSV-A
bootstrap
75
81.25
87.5
93.75
100
B.
CP
Tree scale: 1
SLRSV-C
MH237606.1|Lilium 5875017
MF796980.1|Lilium 14-001
MF796982.1|Lilium 14-002
MF797008.1|Lilium 14-034
MF797010.1|Lilium 14-035
SLRSV-D
OM311160.1|Lily China
MG062674.1|Lilium 17-007
SLRSV-B
MF796998.1|Fragaria 14-023
MZ291960.1|SLRSV Canada
OR513895.1|P.persica France
MF797012.1|Prunus 15-017
NODE 19 length 3526 Carob11-RNA2
NODE 18 length 3541 Carob11-RNA2
NODE 9 length 3268 Carob7-RNA2
NODE 7 length 3505 Carob6-RNA2
NODE 6 length 3543 Carob6-RNA2
NODE 4 length 3446 Carob8-RNA2
NODE 25 length 3322 Carob11-RNA2
NODE 21 length 3450 Carob11-RNA2
NODE 7 length 3548 Carob4-RNA2
NODE 9 length 3452 Carob9-RNA2
NODE 7 length 3400 carob7-RNA2
NODE 6 length 3574 Carob4-RNA2
NODE 7 length 3405 Carob12-RNA2
NODE 5 length 3251 Carob8-RNA2
NODE 20 length 3491 Carob11-RNA2
NODE 6 length 3431 carob12-RNA2
NODE 5 length 3510 Carob12-RNA2
NODE 5 length 3417 carob13-RNA2
SLRSV-E
MZ405641.1|SLRSV Scotland
NC 006965.1|Mentha
MF796996.1|Rubus 14-022
MF796994.1|Clematis 14-021
MF797000.1|Phaseolus 14-024
HM591199.1|Tibouchina-NZ
HM591195.1|Anemone-NZ
HM591197.1|Pepino-NZ
MW854292.1|S.muricatum New Zealand
MF797014.1|Rubus 15-018
MF796978.1|Lilium 13-024
MF796986.1|Lilium 14-008
MF796992.1|Lilium 14-019
MF797006.1|Robinia 14-027
MF796990.1|Lilium 14-010-1
MF797004.1|Rosa 14-026
HM591196.1|Impatiens-NZ
HM591198.1|Rubus New Zealand
MF796974.1|Lilium 12-001
MF797002.1|Rosa 14-025
MN082041.1|P.persica New Zealand
MF796988.1|Lilium 14-010-2
MF796976.1|Lilium 13-023
MF796984.1|Lilium 14-007
SLRSV-A
bootstrap
75
81.25
87.5
93.75
100

### Figure S8

A.

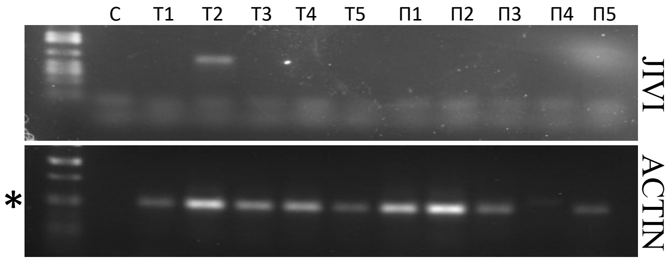


B.

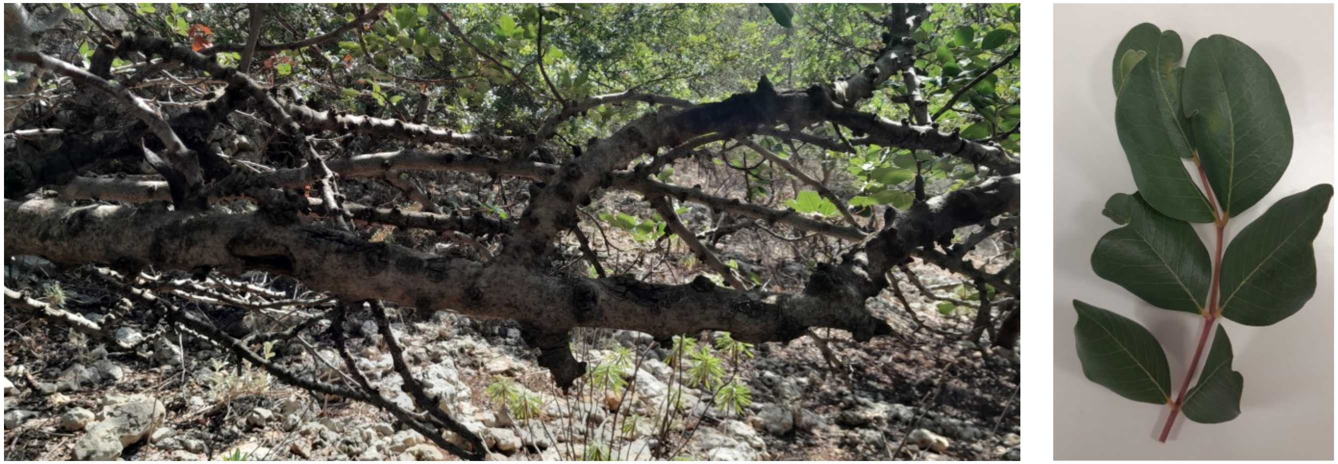

**Figure S9**

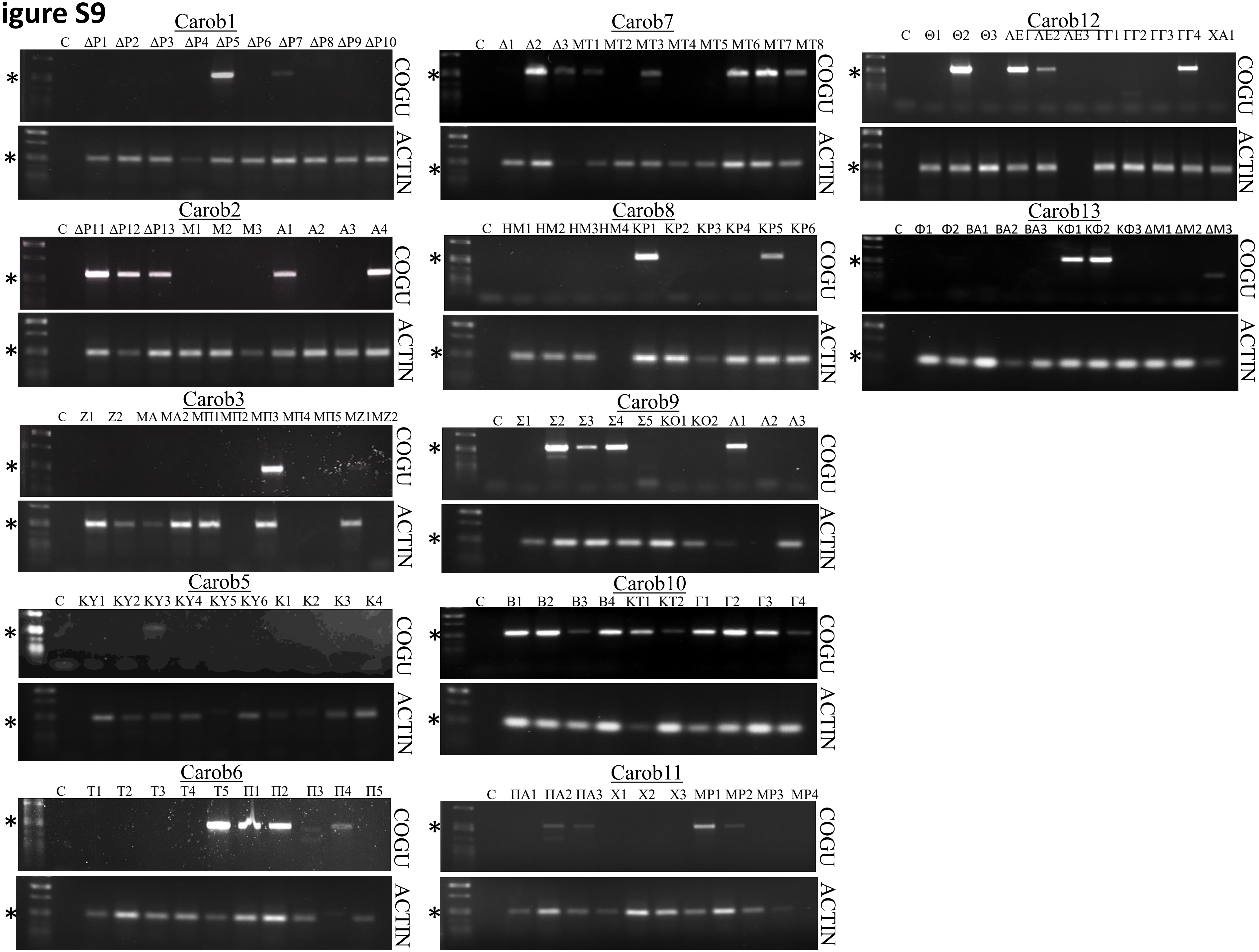

Carob1
C ΔP1 ΔP2 ΔP3 ΔP4 ΔP5 ΔP6 ΔP7 ΔP8 ΔP9 ΔP10
COGU
ACTIN
Carob2
C ΔP11 ΔP12 ΔP13 M1 M2 M3 A1 A2 A3 A4
Carob3
C Z1 Z2 MA MA2 MΠ1 MΠ2 MΠ3 MΠ4 MΠ5 MZ1 MZ2
Carob5
C KY1 KY2 KY3 KY4 KY5 KY6 K1 K2 K3 K4
Carob6
C T1 T2 T3 T4 T5 Π1 Π2 Π3 Π4 Π5
Carob7
C Δ1 Δ2 Δ3 MT1 MT2 MT3 MT4 MT5 MT6 MT7 MT8
Carob8
C HM1 HM2 HM3 HM4 KP1 KP2 KP3 KP4 KP5 KP6
Carob9
C Σ1 Σ2 Σ3 Σ4 Σ5 KO1 KO2 Λ1 Λ2 Λ3
Carob10
C B1 B2 B3 B4 KT1 KT2 Γ1 Γ2 Γ3 Γ4
Carob11
C ΠA1 ΠA2 ΠA3 X1 X2 X3 MP1 MP2 MP3 MP4
Carob12
C Θ1 Θ2 Θ3 ΛE1 ΛE2 ΛE3 ΓΓ1 ΓΓ2 ΓΓ3 ΓΓ4 XA1
Carob13
C Φ1 Φ2 BA1 BA2 BA3 KΦ1 KΦ2 KΦ3 ΔM1 ΔM2 ΔM3

**Figure S10**

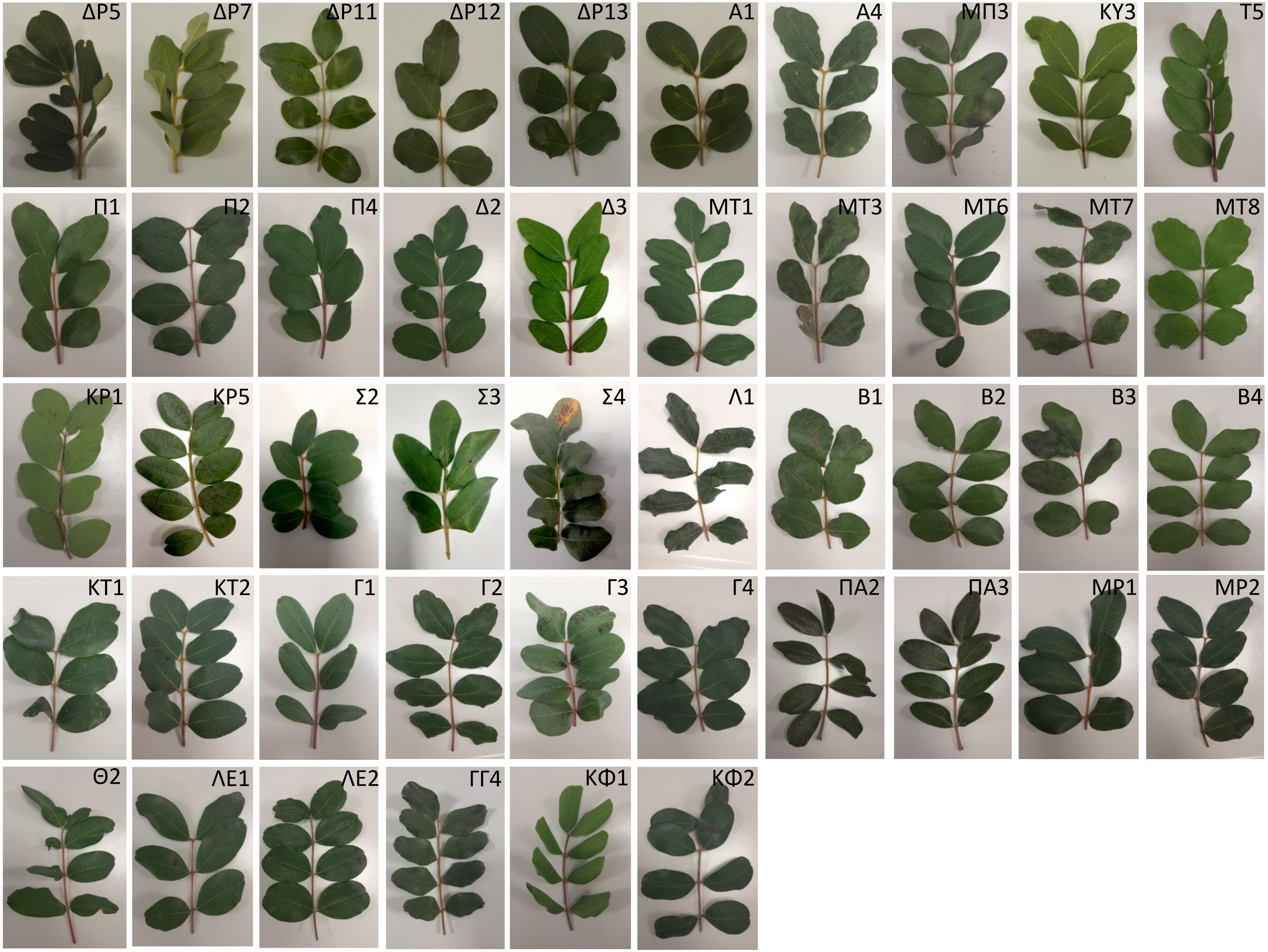
ΔΡ5
ΔΡ7
ΔΡ11
ΔΡ12
ΔΡ13
Α1
Α4
ΜΠ3
ΚΥ3
Τ5
Π1
Π2
Π4
Δ2
Δ3
ΜΤ1
ΜΤ3
ΜΤ6
ΜΤ7
ΜΤ8
ΚΡ1
ΚΡ5
Σ2
Σ3
Σ4
Λ1
Β1
Β2
Β3
Β4
ΚΤ1
ΚΤ2
Γ1
Γ2
Γ3
Γ4
ΠΑ2
ΠΑ3
ΜΡ1
ΜΡ2
Θ2
ΛΕ1
ΛΕ2
ΓΓ4
ΚΦ1
ΚΦ2

**Figure S11**

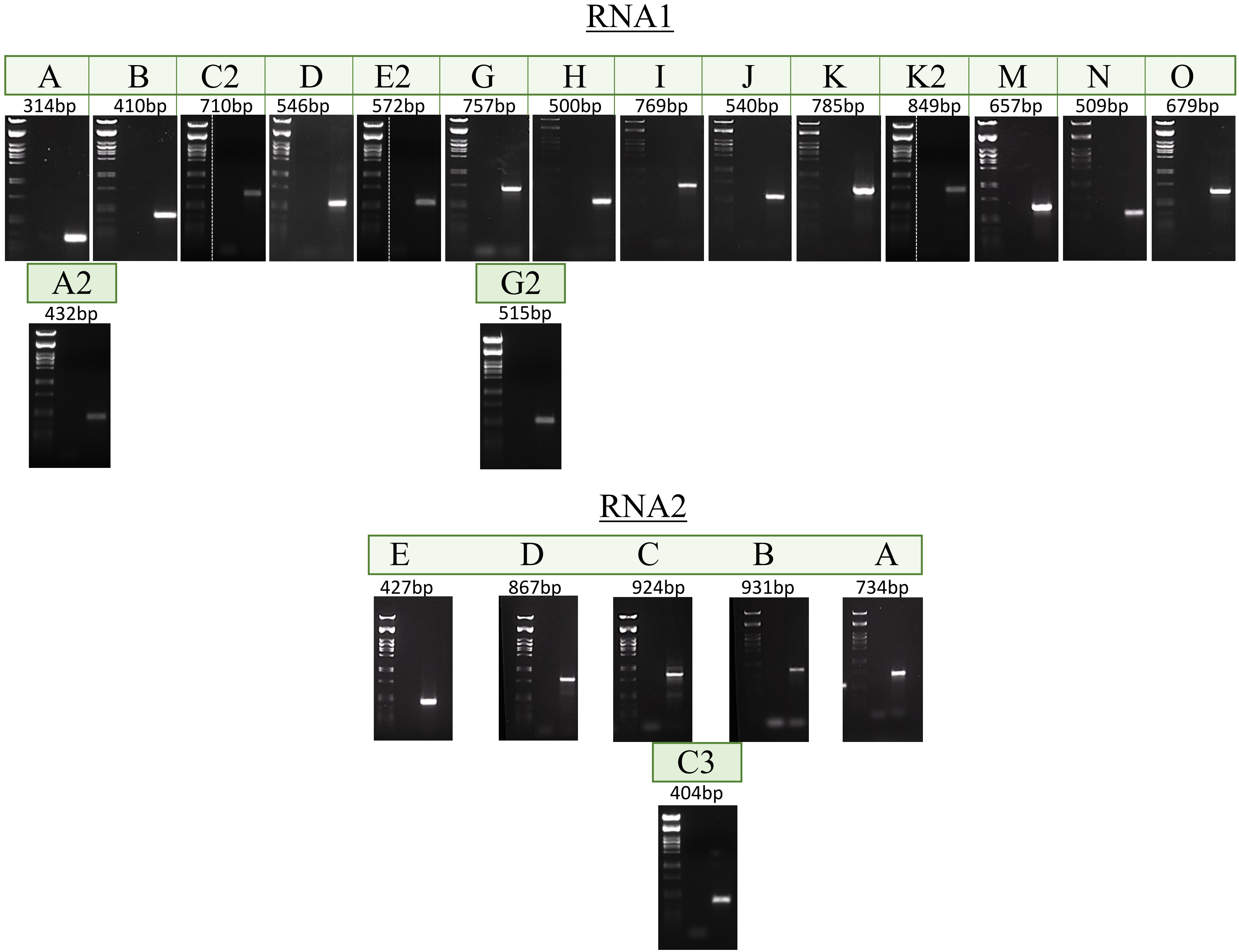

RNA1
A
314bp
B
410bp
C2
710bp
D
546bp
E2
572bp
G
757bp
H
500bp
I
769bp
J
540bp
K
785bp
K2
849bp
M
657bp
N
509bp
O
679bp
A2
432bp
G2
515bp
RNA2
E
427bp
D
867bp
C
924bp
B
931bp
A
734bp
C3
404bp

### Figure S12

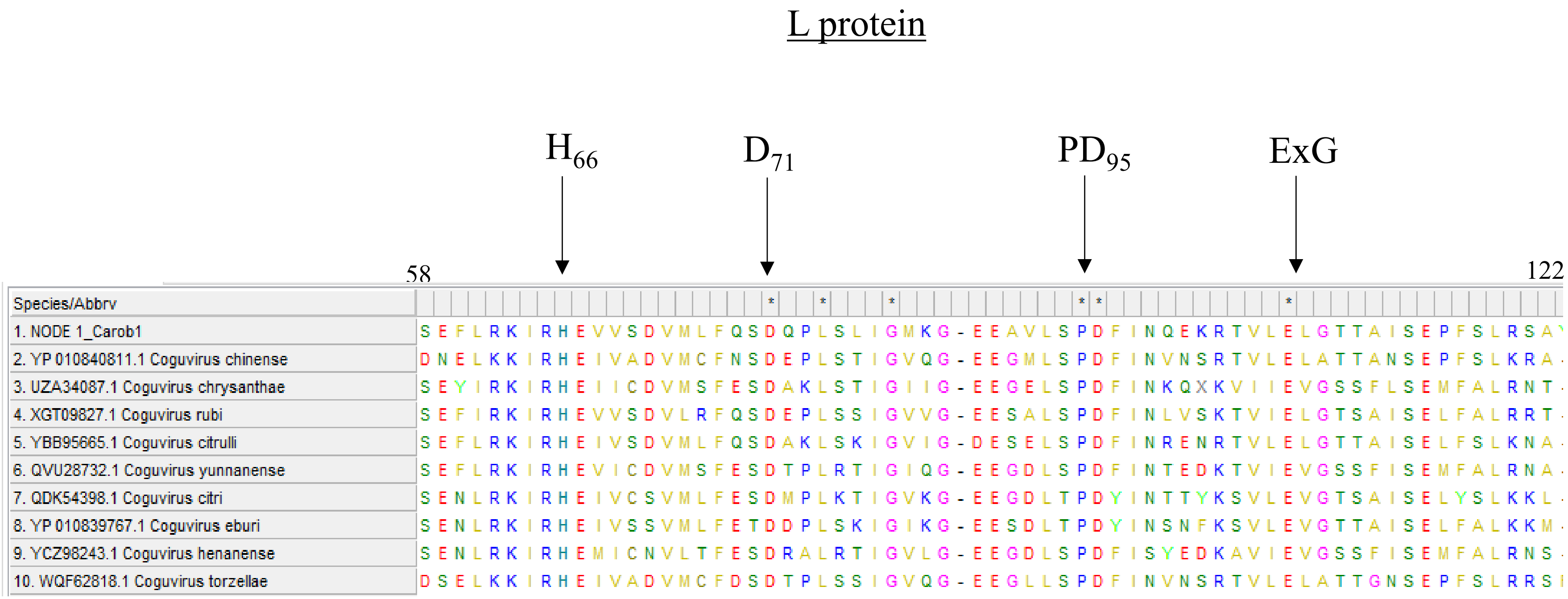
L protein
H66
D71
PD95
ExG
58
122
Species/Abbrv
1. NODE 1_Carob1 SEFLRKIRHEVVSDVMLFQSDQPLSLIGMKG-EEAVLSPDFINQEKRTVLELGTTAISEPFSLRSA
2. YP 010840811.1 Coguvirus chinense DNELKKIRHEIVADVMCFNSDEPLSTIGVQG-EEGMLSPDFINVNSRTVLELATTANSEPFSLKRA-
3. UZA34087.1 Coguvirus chrysanthae SEYIRKIRHEIICDVMSFESDAKLSTIGIIG-EEGELSPDFINKQXKVIIEVGSSFLSEMFALRNT-
4. XGT09827.1 Coguvirus rubi SEFIRKIRHEVVSDVLRFQSDEPLSSIGVVG-EESALSPDFINLVSKTVIELGTSAISELFALRRT-
5. YBB95665.1 Coguvirus citrulli SEFLRKIRHEIVSDVMLFQSDAKLSKIGVIG-DESELSPDFINRENRTVLELGTTAISELFSLKNA-
6. QVU28732.1 Coguvirus yunnanense SEFLRKIRHEVICDVMSFESDTPLRTIGIQG-EEGDLSPDFINTEDKTVIEVGSSFISEMFALRNA-
7. QDK54398.1 Coguvirus citri SENLRKIRHEIVCSVMLFESDMPLKTIGVKG-EEGDLTPDYINTTYKSVLEVGTSAISELYSLKKL-
8. YP 010839767.1 Coguvirus eburi SENLRKIRHEIVSSVMLFETDDPLSKIGIKG-EESDLTPDYINSNFKSVLEVGTTAISELFALKKM-
9. YCZ98243.1 Coguvirus henanense SENLRKIRHEMICNVLTFESDRALRTIGVLG-EEGDLSPDFISYEDKAVIEVGSSFISEMFALRNS-
10. WQF62818.1 Coguvirus torzellae DSELKKIRHEIVADVMCFDSDTPLSSIGVQG-EEGLLSPDFINVNSRTVLELATTGNSEPFSLRRSI

### Figure S13

#### Motif 1

70–82

| Species/Abbrv | | | * | | | * | * | * | | * | | | |
|---|---|---|---|---|---|---|---|---|---|---|---|---|---|
| 1. MP-Carob_1 | P | I | G | T | K | F | Q | Y | A | A | L | T | G |
| 2. YP 010840812.1_Coguvirus chinense | E | P | G | N | R | F | Q | Y | A | A | I | T | C |
| 3. UZA34088.1 Coguvirus chrysanthae | E | E | G | N | R | F | Q | Y | V | A | L | S | G |
| 4. XGT09831.1 Coguvirus rubi | T | E | G | S | R | F | Q | Y | V | A | L | S | G |
| 5. YCZ98241.1 Coguvirus citrulli | S | E | G | S | R | F | Q | Y | V | A | L | T | A |
| 6. YP 010840808.1 Coguvirus yunnanense | E | E | G | N | R | F | Q | Y | V | A | L | S | G |
| 7. XJO56359.1 Coguvirus citri | T | E | G | T | R | F | Q | Y | V | A | L | S | G |
| 8. YEF05680.1 Coguvirus eburi | S | E | G | T | R | F | Q | Y | V | A | L | S | G |
| 9. YCZ98244.1 Coguvirus henanense | E | E | G | K | R | F | Q | Y | V | A | L | V | G |
| 10. WQF62819.1_Coguvirus_tarzelae | V | P | G | S | R | F | Q | Y | A | A | I | T | C |

#### Motif 2

177–185

| Species/Abbrv | | | | * | | * | | * | |
|---|---|---|---|---|---|---|---|---|---|
| 1. MP-Carob_1 | K | V | S | L | H | D | L | R | L |
| 2. YP 010840812.1_Coguvirus chinense | K | V | T | L | H | D | K | R | K |
| 3. UZA34088.1 Coguvirus chrysanthae | S | L | T | L | H | D | T | R | K |
| 4. XGT09831.1 Coguvirus rubi | I | M | S | L | H | D | S | R | R |
| 5. YCZ98241.1 Coguvirus citrulli | I | V | S | L | H | D | S | R | M |
| 6. YP 010840808.1 Coguvirus yunnanense | S | V | T | L | H | D | T | R | K |
| 7. XJO56359.1 Coguvirus citri | S | I | S | L | H | D | S | R | L |
| 8. YEF05680.1 Coguvirus eburi | G | M | S | L | H | D | S | R | L |
| 9. YCZ98244.1 Coguvirus henanense | C | V | D | L | T | D | I | R | K |
| 10. WQF62819.1_Coguvirus_tarzelae | K | V | T | L | H | D | K | R | K |

#### Motif 3

182–194

| Species/Abbrv | * | * | * | | | | * | | * | * | * | | |
|---|---|---|---|---|---|---|---|---|---|---|---|---|---|
| 1. MP-Carob_1 | N | S | N | I | E | N | K | F | E | L | S | T | D |
| 2. YP 010840812.1_Coguvirus chinense | N | S | N | L | S | F | K | F | E | L | S | M | D |
| 3. UZA34088.1 Coguvirus chrysanthae | N | S | N | L | Q | N | K | V | E | L | S | L | D |
| 4. XGT09831.1 Coguvirus rubi | N | S | N | I | S | Q | K | I | E | L | S | L | D |
| 5. YCZ98241.1 Coguvirus citrulli | N | S | N | I | P | Q | K | F | E | L | S | L | D |
| 6. YP 010840808.1 Coguvirus yunnanense | N | S | N | L | Q | N | K | F | E | L | S | M | E |
| 7. XJO56359.1 Coguvirus citri | N | S | N | I | T | Q | K | V | E | L | S | L | D |
| 8. YEF05680.1 Coguvirus eburi | N | S | N | I | T | Q | K | L | E | L | S | L | D |
| 9. YCZ98244.1 Coguvirus henanense | N | S | N | V | P | E | K | F | E | L | S | L | D |
| 10. WQF62819.1_Coguvirus_tarzelae | N | S | N | L | S | F | K | F | E | L | S | M | D |

### Figure S14

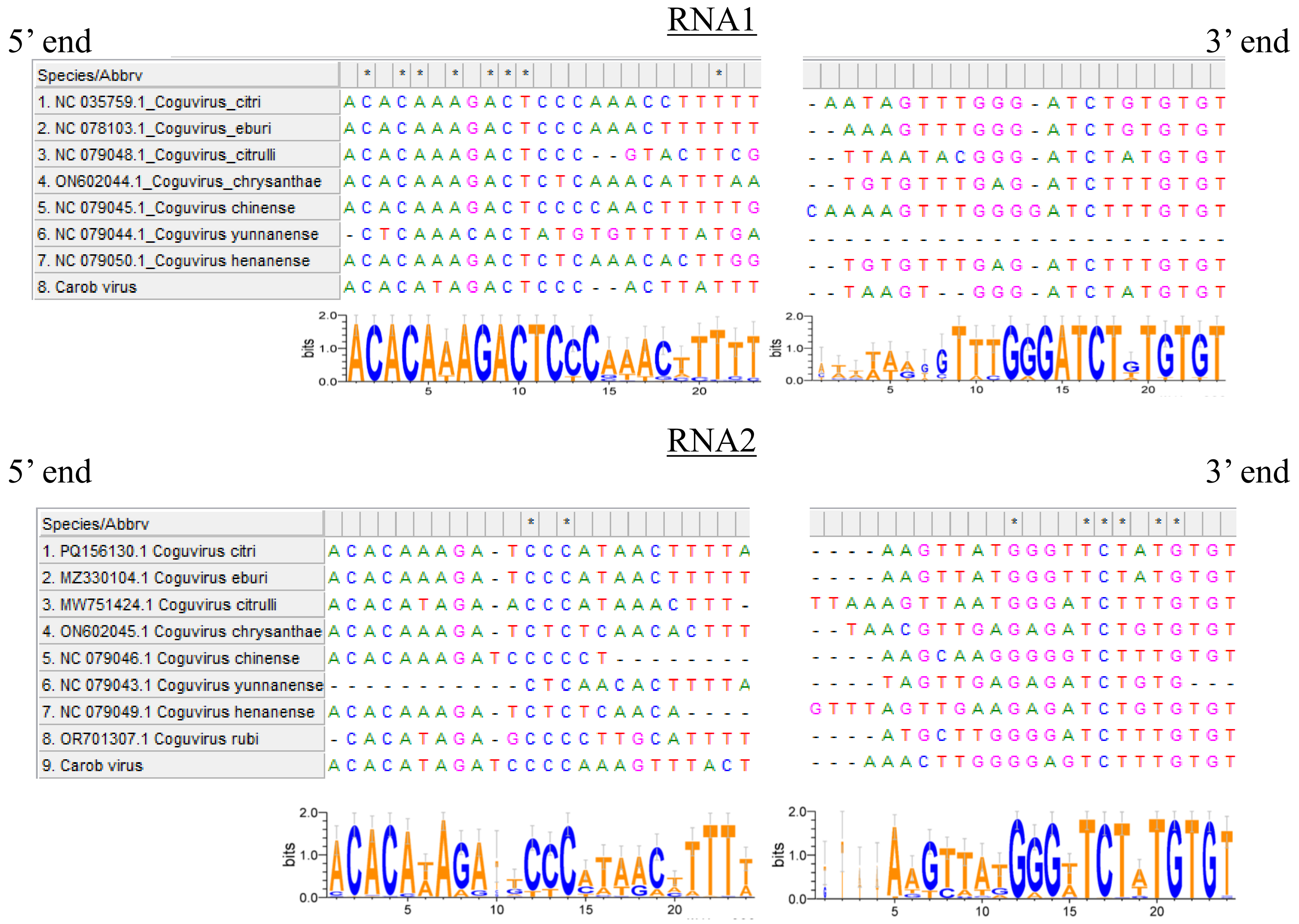

RNA1
5' end
3' end
Species/Abbrv
1. NC 035759.1_Coguvirus_citri
2. NC 078103.1_Coguvirus_eburi
3. NC 079048.1_Coguvirus_citrulli
4. ON602044.1_Coguvirus_chrysanthae
5. NC 079045.1_Coguvirus chinense
6. NC 079044.1_Coguvirus yunnanense
7. NC 079050.1_Coguvirus henanense
8. Carob virus
ACACAAAGACTCCCAAACCTTTTT
ACACAAAGACTCCCAAACTTTTTT
ACACAAAGACTCCC--GTACTTCG
ACACAAAGACTCTCAAACATTTAA
ACACAAAGACTCCCCAACTTTTTG
-CTCAAACACTATGTGTTTTATGA
ACACAAAGACTCTCAAACACTTGG
ACACATAGACTCCC--ACTTATTT
-AATAGTTTGGG-ATCTGTGTGT
--AAAGTTTGGG-ATCTGTGTGT
--TTAATACGGG-ATCTATGTGT
--TGTGTTTGAG-ATCTTTGTGT
CAAAAGTTTGGGGATCTTTGTGT
-----------------------
--TGTGTTTGAG-ATCTTTGTGT
--TAAGT--GGG-ATCTATGTGT
bits
2.0
1.0
0.0
5
10
15
20
RNA2
5' end
3' end
Species/Abbrv
1. PQ156130.1 Coguvirus citri
2. MZ330104.1 Coguvirus eburi
3. MW751424.1 Coguvirus citrulli
4. ON602045.1 Coguvirus chrysanthae
5. NC 079046.1 Coguvirus chinense
6. NC 079043.1 Coguvirus yunnanense
7. NC 079049.1 Coguvirus henanense
8. OR701307.1 Coguvirus rubi
9. Carob virus
ACACAAAGA-TCCCATAACTTTTA
ACACAAAGA-TCCCATAACTTTTT
ACACATAGA-ACCCATAAACTTT-
ACACAAAGA-TCTCTCAACACTTT
ACACAAAGATCCCCCT--------
-----------CTCAACACTTTTA
ACACAAAGA-TCTCTCAACA----
-CACATAGA-GCCCCTTGCATTTT
ACACATAGATCCCCAAAGTTTACT
----AAGTTATGGGTTCTATGTGT
----AAGTTATGGGTTCTATGTGT
TTAAAGTTAATGGGATCTTTGTGT
--TAACGTTGAGAGATCTGTGTGT
----AAGCAAGGGGGTCTTTGTGT
----TAGTTGAGAGATCTGTG---
GTTTAGTTGAAGAGATCTGTGTGT
----ATGCTTGGGGATCTTTGTGT
---AAACTTGGGGAGTCTTTGTGT
bits
2.0
1.0
0.0
5
10
15
20

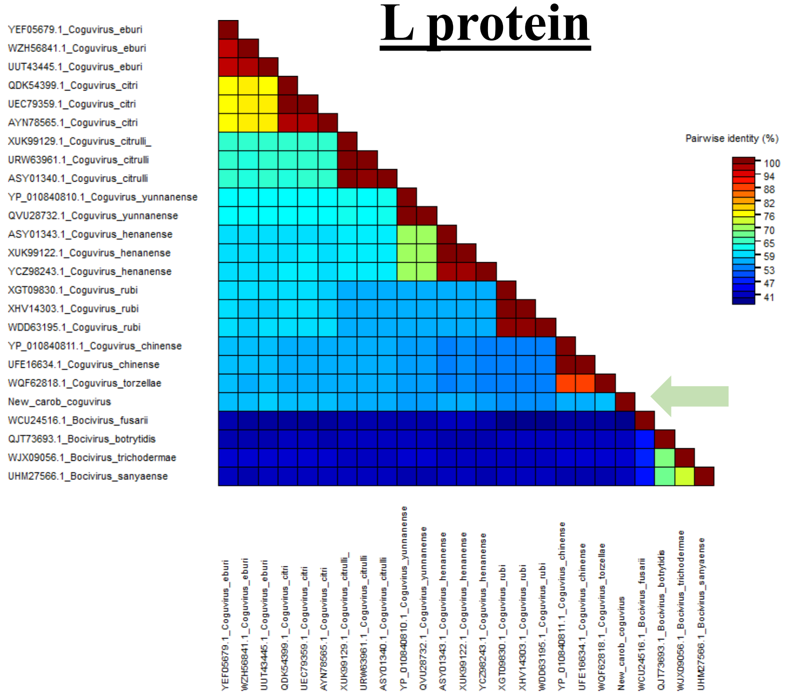
L protein
YEF05679.1_Coguvirus_eburi
WZH56841.1_Coguvirus_eburi
UUT43445.1_Coguvirus_eburi
QDK54399.1_Coguvirus_citri
UEC79359.1_Coguvirus_citri
AYN78565.1_Coguvirus_citri
XUK99129.1_Coguvirus_citrulli_
URW63961.1_Coguvirus_citrulli
ASY01340.1_Coguvirus_citrulli
YP_010840810.1_Coguvirus_yunnanense
QVU28732.1_Coguvirus_yunnanense
ASY01343.1_Coguvirus_henanense
XUK99122.1_Coguvirus_henanense
YCZ98243.1_Coguvirus_henanense
XGT09830.1_Coguvirus_rubi
XHV14303.1_Coguvirus_rubi
WDD63195.1_Coguvirus_rubi
YP_010840811.1_Coguvirus_chinense
UFE16634.1_Coguvirus_chinense
WQF62818.1_Coguvirus_torzellae
New_carob_coguvirus
WCU24516.1_Bocivirus_fusarii
QJT73693.1_Bocivirus_botrytidis
WJX09056.1_Bocivirus_trichodermae
UHM27566.1_Bocivirus_sanyaense
Pairwise identity (%)
100
94
88
82
76
70
65
59
53
47
41

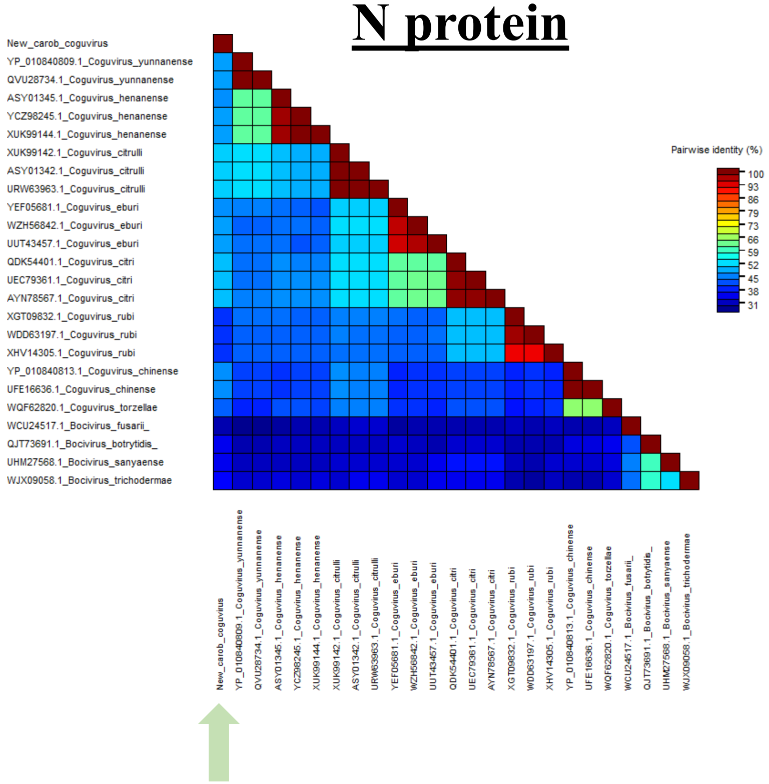
N protein
New_carob_coguvirus
YP_010840809.1_Coguvirus_yunnanense
QVU28734.1_Coguvirus_yunnanense
ASY01345.1_Coguvirus_henanense
YCZ98245.1_Coguvirus_henanense
XUK99144.1_Coguvirus_henanense
XUK99142.1_Coguvirus_citrulli
ASY01342.1_Coguvirus_citrulli
URW63963.1_Coguvirus_citrulli
YEF05681.1_Coguvirus_eburi
WZH56842.1_Coguvirus_eburi
UUT43457.1_Coguvirus_eburi
QDK54401.1_Coguvirus_citri
UEC79361.1_Coguvirus_citri
AYN78567.1_Coguvirus_citri
XGT09832.1_Coguvirus_rubi
WDD63197.1_Coguvirus_rubi
XHV14305.1_Coguvirus_rubi
YP_010840813.1_Coguvirus_chinense
UFE16636.1_Coguvirus_chinense
WQF62820.1_Coguvirus_torzellae
WCU24517.1_Bocivirus_fusarii_
QJT73691.1_Bocivirus_botrytidis_
UHM27568.1_Bocivirus_sanyaense
WJX09058.1_Bocivirus_trichodermae
Pairwise identity (%)
100
93
86
79
73
66
59
52
45
38
31

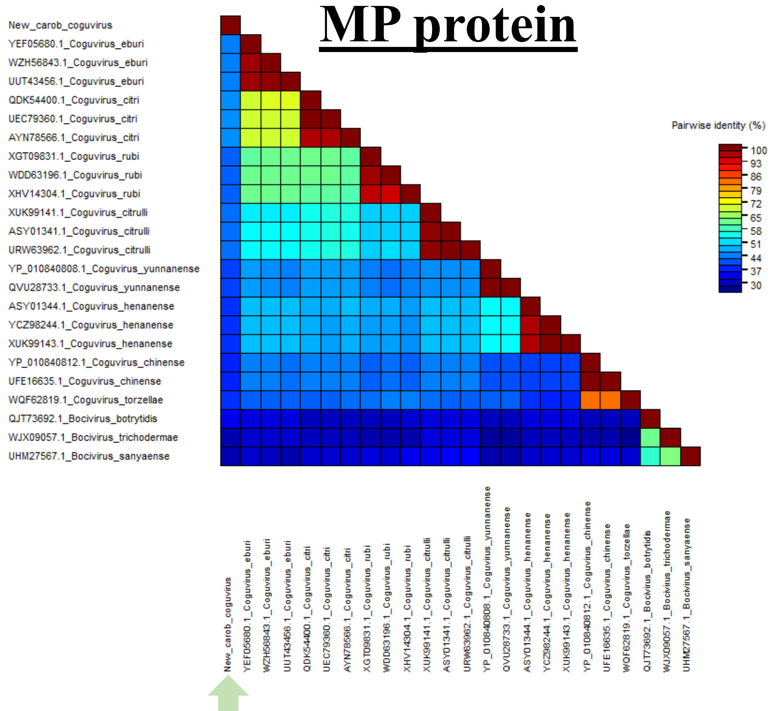
MP protein
New_carob_coguvirus
YEF05680.1_Coguvirus_eburi
WZH56843.1_Coguvirus_eburi
UUT43456.1_Coguvirus_eburi
QDK54400.1_Coguvirus_citri
UEC79360.1_Coguvirus_citri
AYN78566.1_Coguvirus_citri
XGT09831.1_Coguvirus_rubi
WDD63196.1_Coguvirus_rubi
XHV14304.1_Coguvirus_rubi
XUK99141.1_Coguvirus_citrulli
ASY01341.1_Coguvirus_citrulli
URW63962.1_Coguvirus_citrulli
YP_010840808.1_Coguvirus_yunnanense
QVU28733.1_Coguvirus_yunnanense
ASY01344.1_Coguvirus_henanense
YCZ98244.1_Coguvirus_henanense
XUK99143.1_Coguvirus_henanense
YP_010840812.1_Coguvirus_chinense
UFE16635.1_Coguvirus_chinense
WQF62819.1_Coguvirus_torzellae
QJT73692.1_Bocivirus_botrytidis
WJX09057.1_Bocivirus_trichodermae
UHM27567.1_Bocivirus_sanyaense
Pairwise identity (%)
100
93
86
79
72
65
58
51
44
37
30

**Figure S15**
